# Systematic Detection of Large-Scale Multi-Gene Horizontal Transfer in Prokaryotes

**DOI:** 10.1101/2020.08.27.270926

**Authors:** Lina Kloub, Sean Gosselin, Matthew Fullmer, Joerg Graf, J. Peter Gogarten, Mukul S. Bansal

## Abstract

Horizontal gene transfer (HGT) is central to prokaryotic evolution. However, little is known about the “scale” of individual HGT events. In this work, we introduce the first computational framework to help answer the following fundamental question: How often does more than one gene get horizontally transferred in a single HGT event? Our method, called *HoMer*, uses phylogenetic reconciliation to infer single-gene HGT events across a given set of species/strains, employs several techniques to account for inference error and uncertainty, combines that information with gene order information from extant genomes, and uses statistical analysis to identify candidate horizontal multi-gene transfers (HMGTs) in both extant and ancestral species/strains. HoMer is highly scalable and can be easily used to infer HMGTs across hundreds of genomes.

We apply HoMer to a genome-scale dataset of over 22000 gene families from 103 *Aeromonas* genomes and identify a large number of plausible HMGTs of various scales at both small and large phylogenetic distances. Analysis of these HMGTs reveals interesting relationships between gene function, phylogenetic distance, and frequency of multi-gene transfer. Among other insights, we find that (i) the relative frequency of HMGT increases as divergence between genomes increases, (ii) HMGTs often have conserved gene functions, and (iii) rare genes are frequently acquired through HMGT. We also analyze in detail HMGTs involving the zonula occludens toxin and type III secretion systems. By enabling the systematic inference of HMGTs on a large scale, HoMer will facilitate a more accurate and more complete understanding of HGT and microbial evolution.

## Introduction

The transfer of genetic information between organisms that are not in a direct ancestor-descendant relationship, called *Horizontal Gene Transfer* (*HGT*), is a crucial process in microbial evolution. For instance, HGT of pathogenicity and other genomic islands facilitate adaptation to new ecological niches (Hacker et al. 1997, Gogarten et al. 2002, Dobrindt et al. 2004, Papke and Gogarten 2012); HGT helps maintain cohesion within groups or phylotypes of organisms (Papke et al. 2004, Polz et al. 2013); gene transfer, not autochtonous gene duplication, is the most important process for gene family expansion in bacteria and archaea (Treangen and Rocha 2011); and gene transfer together with vertical inheritance shaped the microbial tree of life (Hilario and Gogarten 1993, Doolittle 1999, Andam and Gogarten 2011, Pace et al. 2012). In fact, HGT is so common that the number of distinct genes present in a species far exceeds the number of genes present in any individual genome (Lapierre and Gogarten 2009, Puigbo et al. 2014, Soucy et al. 2015, Fullmer et al. 2015); for example, less then 10% of the non-overlapping gene set from 61 *Escherichia coli* is present in all the genomes that were included in the analysis (Lukjancenko et al. 2010).

Despite the importance of HGT to microbial evolution, surprisingly little is known about the scale of individual HGT events. Specifically, an HGT event may involve the transfer of a gene fragment, a single complete gene, or multiple complete genes, and very little is currently known about the units of HGT events. Chan et al. (2009a) were among the first to conduct a systematic study of the scale of HGT events. The study considered gene families from 144 prokaryotic species and distinguished between HGTs that transferred a complete gene and those that transferred only a part of gene based on finding recombination breakpoints in gene family alignments. The study found that both gene level and sub-gene level HGTs were common and that pathogens were more likely to engage in gene level HGT than non-pathogens. However, this study only considered single-copy gene families and did not study transfers involving multiple genes. A related study by Chan et al. (2009b), using the same methods as Chan et al. 2009a, rejected the hypothesis that protein domains acted as units of HGT. Szöllősi et al. (2015) studied single-gene HGT among fungi and cyanobacteria and, based on gene order information for terminal taxa, they observed that many HGTs between terminal branches appeared clustered together on genomes, suggesting the presence of multi-gene transfers. Phylogenetic analysis coupled with either sequence similarity analysis or phylogenetic reconciliation techniques have also been used to identify some instances of plasmid-borne horizontal transfer of gene clusters (Petersen and Wagner-Dobler 2017, Brinkmann et al. 2018). More recently, Dunning et al. (2019) used multiple grass genomes and phylogenetic comparative analysis to find 59 single-gene HGTs into *Alloteropsis semialata* that were organized into 23 acquired genome fragments, suggesting horizontal transfer of genomic fragments containing multiple genes. While these previous studies have helped establish the presence of multi-gene horizontal transfers, there do not currently exist any rigorous computational frameworks for systematically detecting and quantifying plausible multi-gene horizontal transfers. Researchers have also previously explored “highways of gene sharing” in microbes (Beiko et al. 2005, Zhaxybayeva et al. 2009, Bansal et al. 2013b). These highways represent pairs of species or species groups that are connected to each other by a multitude of HGT events. Highways result when divergent organisms share an ecological niche and engage in gene transfer for extended periods of time. Highways capture the magnitude of HGT that has occurred between a pair of species or species groups but do not shed light on the units of transfer for individual HGT events.

In this work, we focus on the problem of systematic, automated discovery of high-confidence instances where multiple complete genes were transferred in a single horizontal transfer event; we refer to such horizontal transfers as *horizontal multi-gene transfers* (*HMGTs*). We develop a novel computational framework, called *HoMer* (for “<u>Ho</u>rizontal <u>M</u>ulti-gene transf<u>er</u>”), that builds upon recent computational advances in the detection of single-gene HGTs and leverages large-scale availability of microbial genomic datasets to infer plausible HMGTs. HoMer infers single-gene HGT events across the given set of species or strains using phylogenetic reconciliation, uses several techniques to account for (single-gene) HGT inference uncertainty, combines that information with gene order information, and uses statistical analysis to identify candidate (multi-gene) HMGTs. HoMer can infer HMGTs not only between terminal taxa but also between ancestral species (internal edges) on the species tree, allows for easy adjustment of the stringency of detected HMGTs, and can be used to estimate statistical support for the inferred HMGTs. It is also highly scalable and can be applied to hundreds of taxa in a matter of hours.

We apply HoMer to a genome-scale dataset of over 22000 gene families (or consolidated homologous groups) from 103 *Aeromonas* strains representing 28 different species (Rangel et al. 2019), and infer a large number of plausible HMGTs of various scales at both small and large phylogenetic distances. *Aeromonas* are a genus of gram-negative bacteria that are known to cause disease in humans and fish. They are found in water and sediments and live in beneficial associations with fish and leeches (Janda and Abbott 2010, Fernandez-Bravo and Figueras 2020, Marden et al. 2016, Milligan-Myhre et al. 2011). The *Aeromonas* genus serves as an excellent test case for this study because of the availability of genomes from 28 distinct species with multiple strain genomes available for several of these species, resulting in a broad dataset with sufficient breadth and depth to assess both inter- and intra-species HGTs and HMGTs. Moreover, the presence of frequent HGT within the Aeromonads has been previously established (Silver et al. 2011, Morandi et al. 2005, Colston et al. 2014).

Analysis of HMGTs inferred on the *Aeromonas* dataset reveals several fundamental insights and interesting relationships between gene function, phylogenetic distance, and frequency of multi-gene transfer. For instance, we find that (i) the relative frequency of HMGT increases as divergence between genomes increases, (ii) genes transferred together in an HMGT often belong to the same COG functional category, and (iii) rare genes are frequently acquired through HMGT. We also analyze in detail some specific HMGTs involving type III secretion systems (T3SS) and the zonula occludens toxin (ZOT). Two types of T3SSs were previously described in *Aeromonas* spp. (Bomar et al. 2013, Rangel et al. 2019). We found components of T3SS-1 and components of T3SS-2 to have significantly different phylogenetic reconstructions. Specifically, components of the T3SS-1 apparatus, present in about 50% of the analyzed *Aeromonas* genomes, group with homologs from *Yersinia* in phylogenetic reconstructions, while components of T3SS-2, identified in only six of the analyzed *Aeromonas* genomes, group with the homologs from a *Salmonella enterica* subsp. *enterica* T3SS. The *Aeromonas* ZOTs represent three divergent families. Inspection of the genes surrounding the ZOTs suggests phages as vehicles for these transfers. We find that these pathogenicity related genes are transferred more frequently within genera than between genera. The within-genus transfers and recombination events lead to a diversification of possible symbiotic interactions.

This work makes it feasible, for the first time, to systematically infer HMGTs on a large scale, and demonstrates the prevalence and significance of HMGTs in microbial evolution. The systematic discovery of HMGTs, enabled by HoMer, will help advance our understanding of horizontal gene transfer and microbial evolution.

HoMer is freely available open-source from https://compbio.engr.uconn.edu/software/homer/. The *Aeromonas* dataset used in this work and a complete list of putative HMGTs discovered for this dataset are also freely available from the same URL.

## New Approaches

There are four key challenges in designing a computational framework for systematic, large-scale discovery of HMGTs. First, erroneous HGT inference. Second, unavailability of genomes (specifically, gene orders) for internal/ancestral nodes in the species phylogeny. Third, precisely defining an HMGT. And fourth, controlling the false discovery rate for HMGTs.

Current approaches focus on discovering HGT of single genes and can have high false-positive and falsenegative rates. Our method, HoMer, infers single-gene HGT events across the given set of species, uses several techniques to account for inference uncertainty, combines that information with gene order information, and uses statistical analysis to identify candidate horizontal multi-gene transfers (HMGTs). We briefly describe the key steps in HoMer below.

1. *Inference of high-confidence HGTs*. To infer HGTs we used a recently developed reconciliation-based technique that reconciles gene trees with a given species tree under a model that accounts for gene duplication, gene loss and horizontal gene transfer. In our inference, we account for several sources of HGT inference uncertainty, such as gene tree error, transfer inference uncertainty, and uncertainty of assigning the donor and recipient for a transfer.
2. *Map HGTs to genomic locations*. For each possible donor-recipient pair in the species tree, we map the locations of the inferred HGTs from that donor to that recipient along the donor (and/or recipient) genome using the available gene orders at the leaves of the species tree.
3. *Define HMGTs for transfers between extant species*. We define HMGTs to be regions of the donor and/or recipient genome that have “unusually many” high-confidence HGTs clustered together. We define HMGTs formally using three parameters 〈*x, y, z*〉, where we first identify contiguous regions of *y* genes in which at least *x* genes have been transferred, and then merge the identified regions with neighboring regions or HGTs if the distance between them is no more than z. This is illustrated in Figure 1. For appropriate values of 〈*x,y, z*〉, e.g., 〈3, 4, 1〉, each of these merged regions constitutes a plausible HMGT. In defining these regions we also account for the presence of rare genes that occur very infrequently in the considered species (and which may have been acquired by HGT from an external species after an HMGT event).
4. *Define HMGTs for transfers between ancestral species*. Since gene-orders are only available for extant species, to infer HMGTs between ancestral species we look for HMGT regions using the most compliant ordering of any of the extant descendants of the donor species (and/or recipient species).
5. *Statistical analysis to determine false discovery rate*. To determine the appropriate 〈*x, y, z*〉 values to use for any given dataset, we use simulations where the inferred HGTs are appropriately randomized (preserving total counts as well as donors and recipients) and HMGTs are inferred using these randomized HGTs. This allows for the estimation of the false discovery rate for any specific assignment of 〈*x, y, z*〉 for the given dataset. Further details on these and other aspects of HoMer appear in the Materials and Methods section.

**Figure 1.**
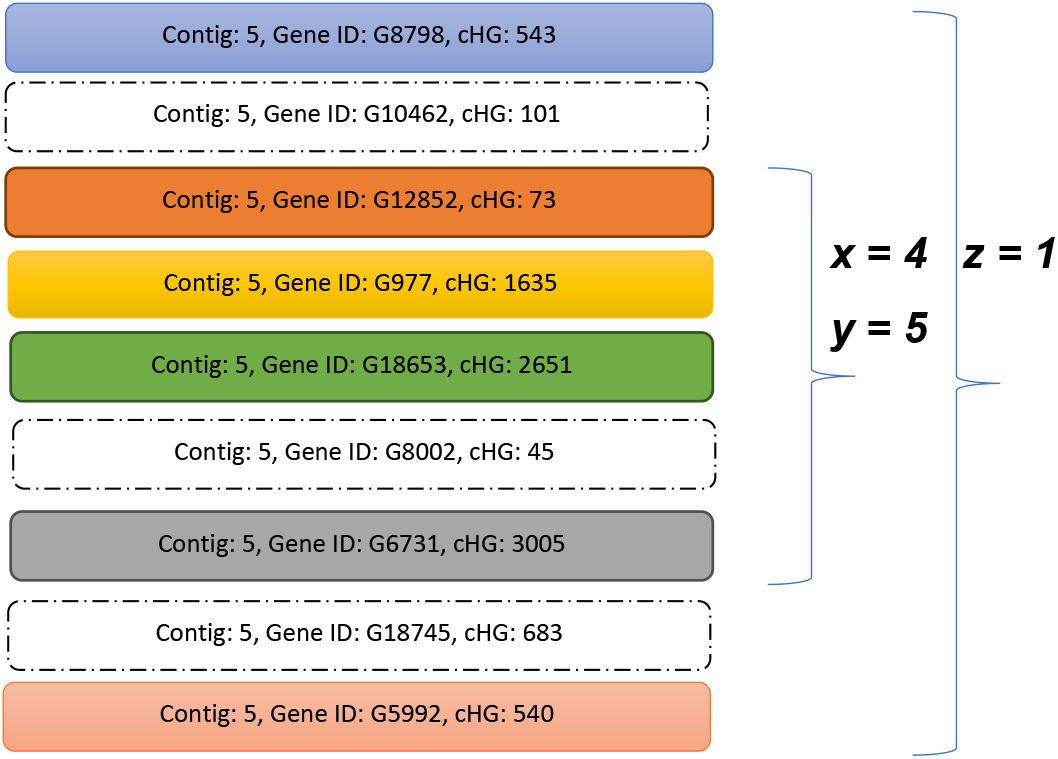
Inferring HMGTs using 〈*x, y, z*〉 parameters. The figure depicts a part of a genome ordering (genes as blocks ordered from top to bottom) along a specific contig from the donor (or recipient) species. The colored (filled) blocks represent genes that were detected as transferred for that donor-recipient pair. With 〈*x,y*〉 = (4,5), the contiguous block consisting of genes G12825 through G6731 would be identified as a transferred region since it consists of 5 genes out of which at least 4 are transferred. Finally, using the region extension parameter *z* = 1, the nearby transferred genes G8798 and G5992 would be merged with the identified transferred region to form a single merged HMGT consisting of all the genes shown in the figure. Note that 〈*x, y*〉 regions can be ambiguous; for example, in this figure genes G8798 through G18653 also form an 〈*x, y*〉 = (4,5) region. However, as long as the region extension parameter *z* is chosen so that *z* ≥ *y* – *x*, the merged HMGTs will be unambiguous.

## Results

We applied HoMer to a genome-scale dataset of 22282 consolidated homologous groups (cHGs), i.e, gene families, from 103 *Aeromonas* genomes. The 103 genomes in this dataset correspond to 28 different species. Of these 28 different species, 18 are represented by a single genome, while the remaining 10 are each represented by at least two genomes corresponding to different strains from that species. This allows us to infer and compare HGT and HMGT patterns for donor-recipient pairs from the same species and from different species. We refer to these two types of HGTs/HMGTs as within-species HGTs/HMGTs and across-species HGTs/HMGTs, respectively.

As described in detail below, our analysis identifies a large number of putative HMGTs both within-species and across-species, and clearly demonstrates that average transfer size is much higher across-species than within species. Contrary to our expectations, we find that there is little difference between the functions associated with HMGT and HGT genes. We also identify frequent HMGTs of rare genes from species not represented in our species tree, and address two specific biological questions; one related to HMGT frequency and phylogenetic distance and the other related to the functions of genes transferred in a single HMGT event. We also perform an in-depth biological analysis of some of the identified HMGTs.

### Terminology

In presenting our results, we use the following terminology.

*Leaf-to-leaf HGT or HMGT*: An HGT or HMGT where the donor and recipient are both leaf-edges on the species tree.
*Internal HGT or HMGT*: Any HGT or HMGT that is not leaf-to-leaf.
*Wthin-species HGT or HMGT*: A leaf-to-leaf HGT or HMGT that occurs between two strains of the same species. *Across-species HGT or HMGT*: A leaf-to-leaf HGT or HMGT that occurs between leaf-edges corresponding to different species.

Note that any HGT or HMGT is either a within-species, across-species, or internal HGT or HMGT.

#### HMGTs are widespread both within and across species

Our analysis identifies a large number of HMGTs from a large number of distinct donor-recipient pairs. Recall that we infer plausible HMGTs using the three parameters 〈*x, y, z*〉, where we first identify contiguous regions of *y* genes in which at least *x* genes have been transferred, and then merge the identified regions with neighboring regions or HGTs if the distance between them is no more than *z*. Using our default, statistically supported setting of 〈3,4,1〉 for these 〈*x, y, z*〉 parameters, we identified 337 plausible within-species HMGTs from 144 distinct donor-recipient pairs, 163 plausible across-species HMGTs from 129 distinct donor-recipient pairs, and 345 plausible internal HMGTs from 141 distinct donor-recipient pairs. These HMGTs contained an average of 3.42 detected high-confidence HGTs. Table 1 shows detailed results for all 〈*x, y, z*〉 parameters settings considered. As the table shows, we find a much larger number of smaller HMGTs (5823 total HMGTs containing an average of 2.22 detected high-confidence HGTs using parameter setting 〈2,3,1〉), as well as a significant number of larger HMGTs (183 total HMGTs containing an average of 4.63 detected high-confidence HGTs using parameter setting 〈4,5,1〉). As Table 1 also shows, these results remain remarkable consistent as the value of *z* is increased from 1 to 2, suggesting that the boundaries of inferred HMGTs are largely accurate.

**Table 1.**
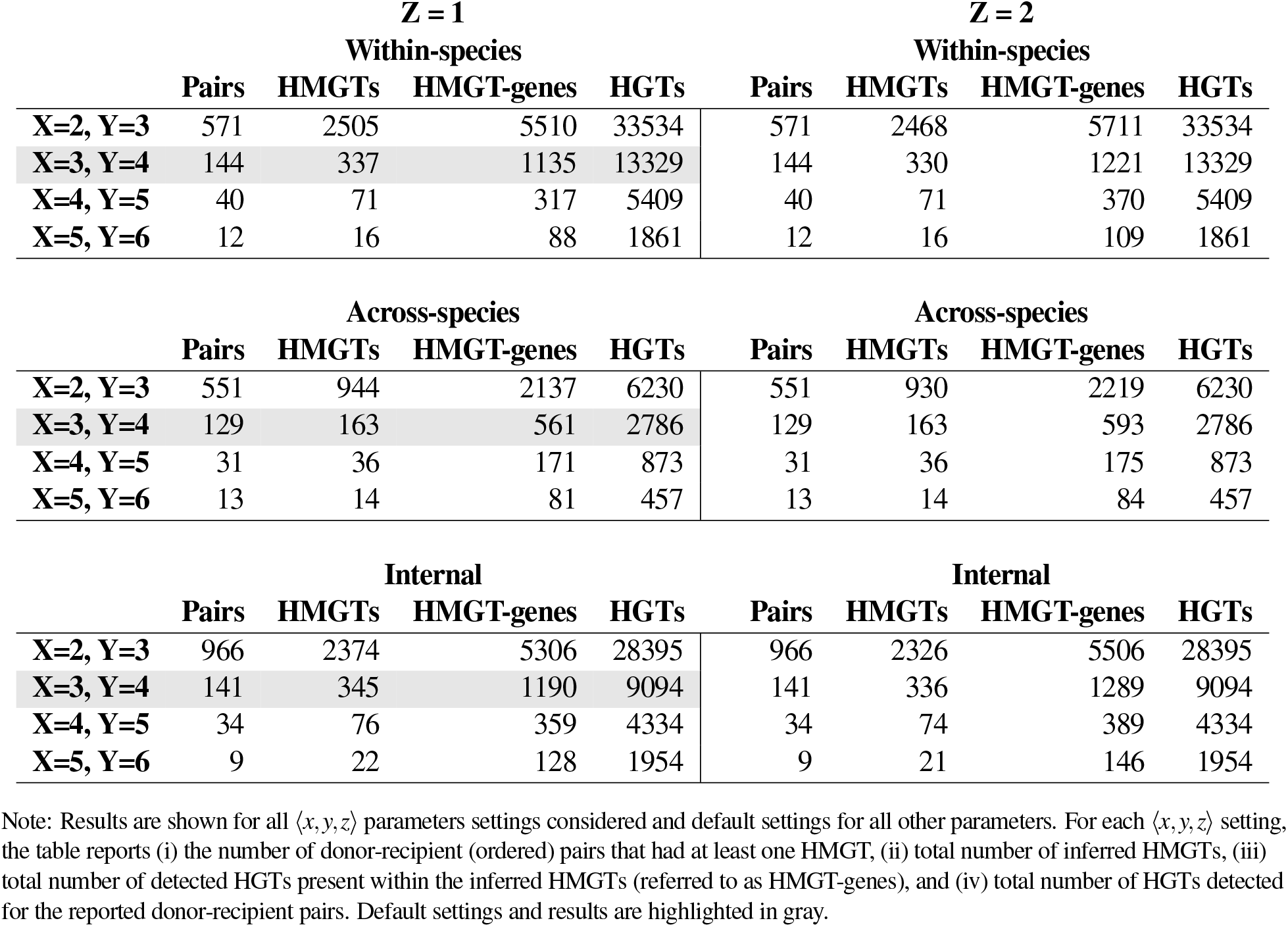
Results of HMGT inference analysis on the *Aeromonas* dataset.

| **Z = 1** |  |  |  |  | **Z = 2** |  |  |  |
| --- | --- | --- | --- | --- | --- | --- | --- | --- |
| **Within-species** |  |  |  |  | **Within-species** |  |  |  |
|  | Pairs | HMGTs | HMGT-genes | HGTs | Pairs | HMGTs | HMGT-genes | HGTs |
| **X=2, Y=3** | 571 | 2505 | 5510 | 33534 | 571 | 2468 | 5711 | 33534 |
| **X=3, Y=4** | 144 | 337 | 1135 | 13329 | 144 | 330 | 1221 | 13329 |
| **X=4, Y=5** | 40 | 71 | 317 | 5409 | 40 | 71 | 370 | 5409 |
| **X=5, Y=6** | 12 | 16 | 88 | 1861 | 12 | 16 | 109 | 1861 |

| **Across-species** |  |  |  |  | **Across-species** |  |  |  |
| --- | --- | --- | --- | --- | --- | --- | --- | --- |
|  | Pairs | HMGTs | HMGT-genes | HGTs | Pairs | HMGTs | HMGT-genes | HGTs |
| **X=2, Y=3** | 551 | 944 | 2137 | 6230 | 551 | 930 | 2219 | 6230 |
| **X=3, Y=4** | 129 | 163 | 561 | 2786 | 129 | 163 | 593 | 2786 |
| **X=4, Y=5** | 31 | 36 | 171 | 873 | 31 | 36 | 175 | 873 |
| **X=5, Y=6** | 13 | 14 | 81 | 457 | 13 | 14 | 84 | 457 |

| **Internal** |  |  |  |  | **Internal** |  |  |  |
| --- | --- | --- | --- | --- | --- | --- | --- | --- |
|  | Pairs | HMGTs | HMGT-genes | HGTs | Pairs | HMGTs | HMGT-genes | HGTs |
| **X=2, Y=3** | 966 | 2374 | 5306 | 28395 | 966 | 2326 | 5506 | 28395 |
| **X=3, Y=4** | 141 | 345 | 1190 | 9094 | 141 | 336 | 1289 | 9094 |
| **X=4, Y=5** | 34 | 76 | 359 | 4334 | 34 | 74 | 389 | 4334 |
| **X=5, Y=6** | 9 | 22 | 128 | 1954 | 9 | 21 | 146 | 1954 |

Note: Results are shown for all $\langle x, y, z \rangle$ parameters settings considered and default settings for all other parameters. For each $\langle x, y, z \rangle$ setting, the table reports (i) the number of donor-recipient (ordered) pairs that had at least one HMGT, (ii) total number of inferred HMGTs, (iii) total number of detected HGTs present within the inferred HMGTs (referred to as HMGT-genes), and (iv) total number of HGTs detected for the reported donor-recipient pairs. Default settings and results are highlighted in gray.

The analysis also shows that both HGTs and HMGTs are far more frequent between within-species donor-recipient pairs than between across-species donor-recipient pairs. For instance, with default parameter settings, we observed an average of 92.56 HGTs and 2.34 HMGTs among the 144 identified within-species donor-recipient pairs, but only an average of 21.6 HGTs and 1.26 HMGTs among the 129 identified across-species donor-recipient pairs. As Table 1 shows, this trend holds across all 〈*x, y, z*〉 parameter settings used in the analysis.

We found little difference between the average sizes of the inferred HMGTs within and across species. Using default parameters, we observed that within-species HMGTs contained an average of 3.37 detected HGTs and across-species HMGTs contained an average of 3.44 detected HGTs. As Table 1 shows, this observation holds across all used 〈*x, y, z*〉 parameter settings. We point out that actual HMGTs sizes (i.e., number of genes transferred in an HMGT event) may be larger than the average sizes reported here since we only count the number of *detected* HGTs present in each HMGT.

Figures 2 and 3 show Circos plots (Krzywinski et al. 2009) displaying all across-species and within-species HMGTs, respectively, inferred using default parameter settings. Corresponding figures displaying HMGT-genes appear in Supplementary Figures S3 and S4.

**Figure 2.**
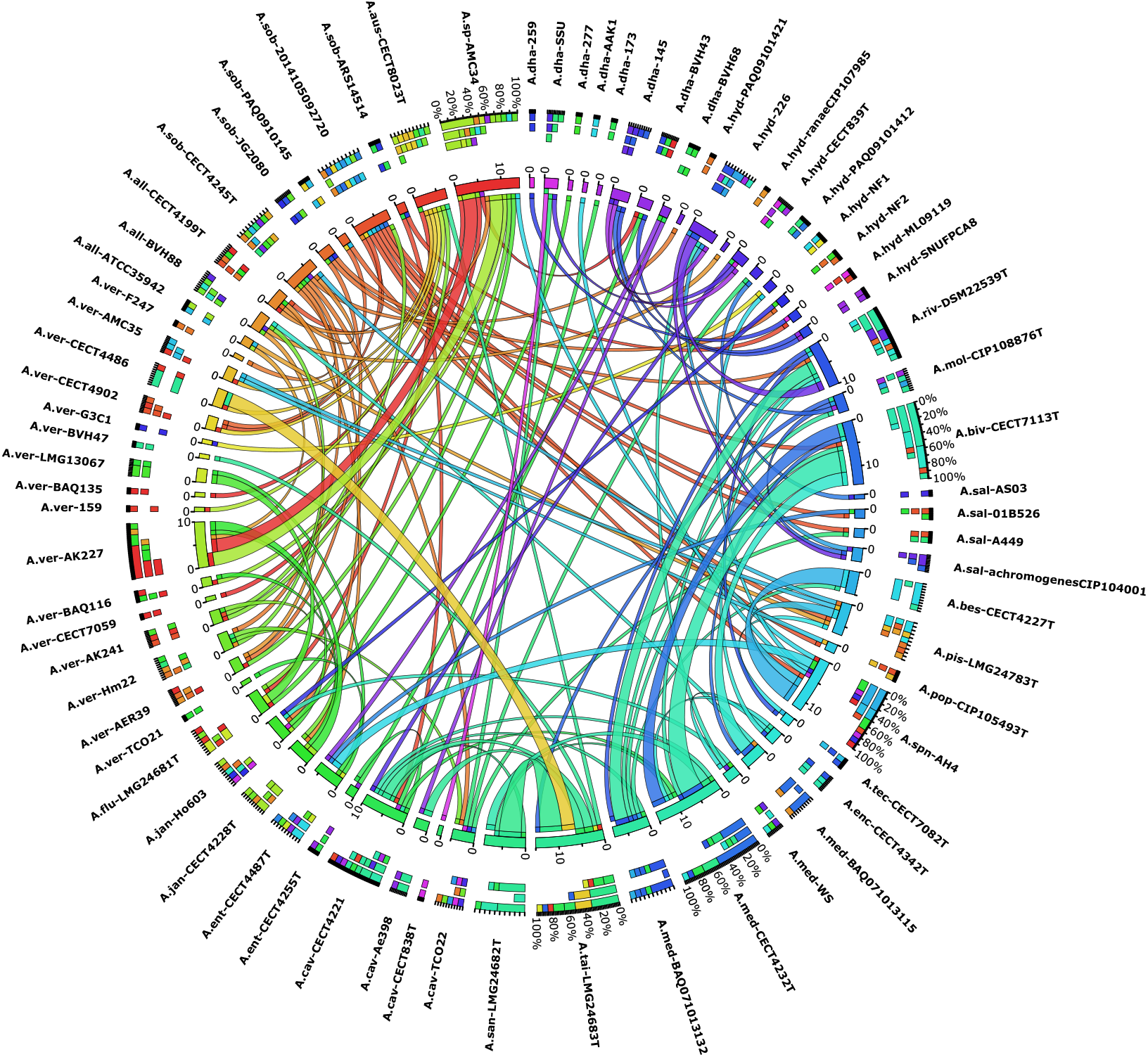
Across-species HMGTs. Each ribbon connects two *Aeromonas* genomes from different species and corresponds to inferred across-species HMGTs between those two genomes. Ribbons are colored according to the color of the donor genome (the color for each genome is shown on the associated segment in the inner ring). The tip of a ribbon at the donor end is colored according to the recipient genome’s color. The thickness of a ribbon corresponds to the number of HMGTs for that donor-recipient pair, as quantified by the numbers around each segment in the inner ring. For each genome, both incoming (where that genome serves as recipient) and outgoing (where that genome serves as donor) ribbons are shown. The outer ring shows three stacked columns for each genome. Among these three stacked columns, the inner column shows the color distribution of recipients for outgoing ribbons, the middle column shows the color distribution of donors for incoming ribbons, and the outer column shown the combined color distribution for both incoming and outgoing ribbons, for that genome. The figure only includes those *Aeromonas* genomes that served as donor or recipient for at least one across-species HMGT. Only HMGTs inferred using default parameters are shown.

**Figure 3.**
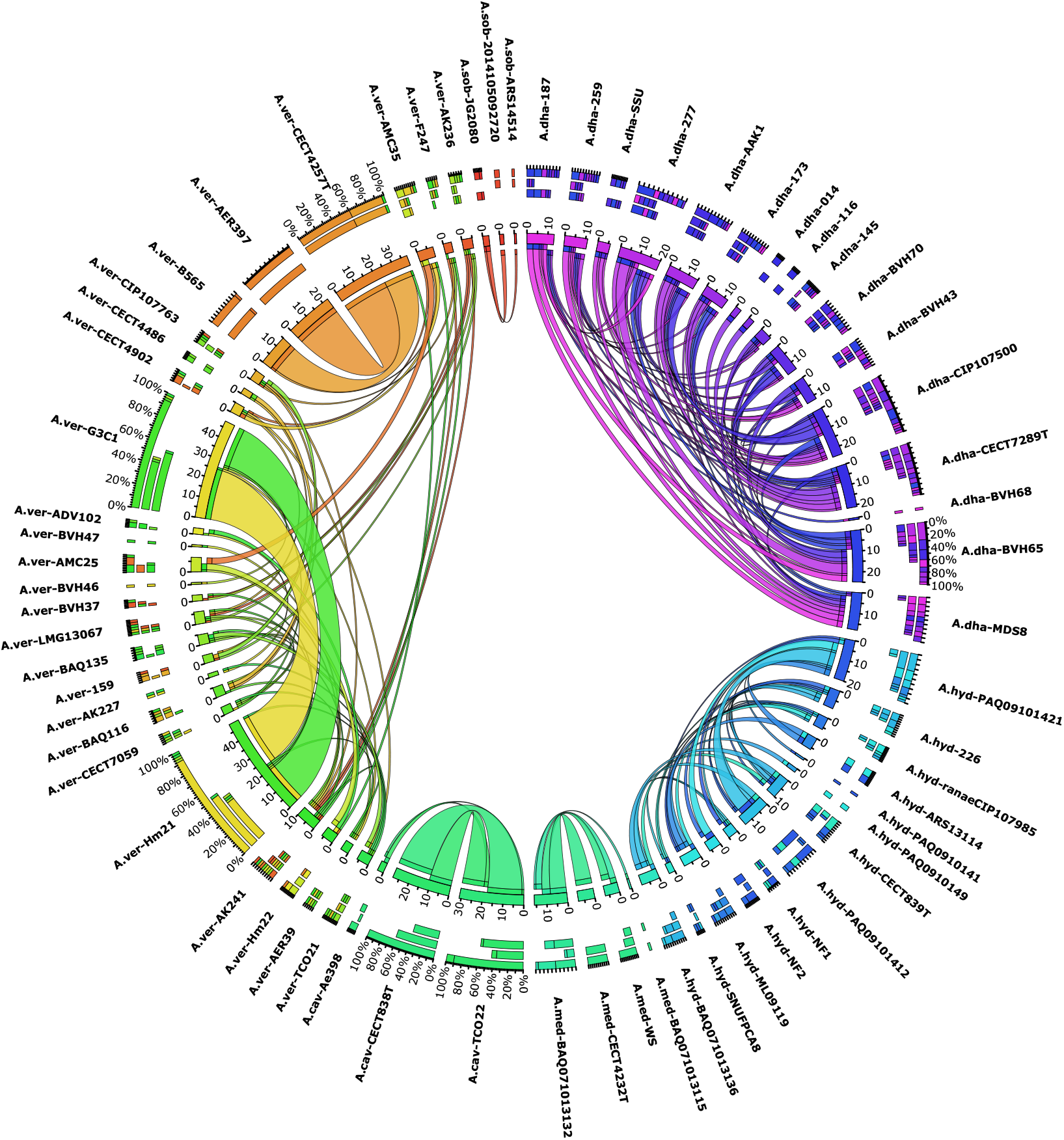
Within-species HMGTs. Each ribbon connects two *Aeromonas* genomes from the same species and corresponds to inferred within-species HMGTs between those two genomes. Interpretation is identical to that of Figure 2. Only HMGTs inferred using default parameters are shown.

It is worth noting that our analysis likely under-reports the number of within-species HGTs and HMGTs by a larger fraction than the number of across-species HGTs and HMGTs. Our HGT detection approach relies on well-supported phylogenetic discordance between gene trees and the species tree. Since within-species genomes are generally very similar, this approach would tend to underestimate the number of within-species HGTs. This, in turn, could result in an underestimation of the number and/or size of inferred HMGTs.

#### HMGT patterns are different within and across species

Despite the similarity in HMGT sizes within- and across-species, we find that the *relative frequency* of HMGT is significantly higher across-species than within-species. Specifically, we find that for across-species donor-recipient pairs a far larger fraction of total HGTs were transferred as part of HMGTs than for within-species donor-recipient pairs. For instance, using default parameter values, we observed that a total of 20.1% of the detected HGTs were contained within HMGTs for the 129 identified across-species donor-recipient pairs, while only a total of 8.5% of the detected HGTs were contained inside HMGTs for the 144 identified within-species donor-recipent pairs. Furthermore, as Figure 4 shows, among the 129 identified across-species donor-recipient pairs, almost half had at least 50% of their detected HGTs contained inside HMGTs, and 9% of the pairs had over 90% of their detected HGTs contained inside HMGTs. This is in stark contrast with within-species donor-recipient pairs, where none of the pairs had at least 50% of their detected HGTs contained inside HMGTs. These results suggest that as divergence between genomes increases, the relative frequency of HMGT increases as well.

**Figure 4.**
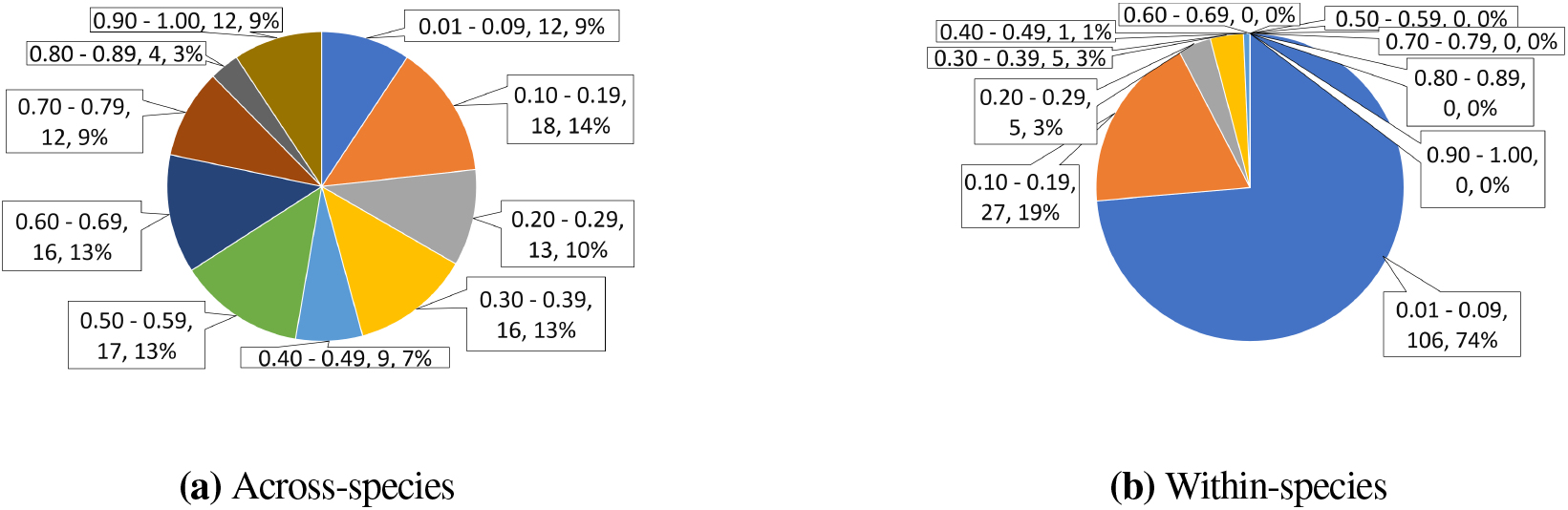
The two pie charts show distributions of the fraction of detected HGTs contained inside HMGTs for the identified across-species donor-recipient pairs (part (a)) and within-species donor-recipient pairs (part (b)). Each slice label consists of three parts; the first part is the range (fraction of detected HGTs contained inside HMGTs) that the slice represents, the second part is the number of donor-recipient pairs that make up that slice, and the third part is the percent area of the pie occupied by that slice.

#### HMGT inference is robust to parameter choices

To assess the robustness of inferred HMGTs and of the observations made above, we evaluated the impact of our specific parameter choices on results. Available parameters and their default settings used in our HMGT inference framework, HoMer, are described in detail in the “Materials and Methods” section and are summarised in Supplementary Table S1. The impact of using different 〈*x, y, z*〉 parameters has already been discussed in detail above (Table 1). Here we discuss the impact of changing other parameter settings on results.

##### More permissive HGT inference

To reduce the number of false-positive HGTs, at the risk of greater false-negative HGTs, we used a high default transfer cost of 4 for HGT inference. We repeated the analysis with a smaller transfer cost of 3, the default recommended cost in the HGT inference method employed (Bansal et al. 2018). These results are shown in Supplementary Table S3. Comparing these numbers with those reported in Table 1, we find that many more HGTs and HMGTs are inferred within-species, and a few more are inferred across-species. Surprisingly, internal HMGT results remain largely unaffected. Despite the higher numbers of within- and across-species HMGTs and HGTs inferred, all observations made above, e.g., those regarding HGT and HMGT abundance and patterns within- and across-species, remain unaffected.

##### More stringent HGT mapping threshold

There can be considerable uncertainty in assigning specific donors and recipients to inferred HGTs. To address this problem, in our analysis, we only use HGTs that have at least 51% support (in the underlying phylogenetic reconciliation) for both the donor mapping and the recipient mapping. This 51% threshold balances the need to infer relatively accurate mappings with the need to not have a very high false-negative rate for usable HGTs. To assess how HMGT inference would be impacted if using a stricter mapping threshold, we repeated the analysis with a mapping threshold of 75% for both donors and recipients. As expected, this results in a very high a false-negative rate for usable HGTs and the numbers of inferred donor-recipient pairs, HMGTs, and HGTs, are all substantially reduced. Supplementary Table S2 shows these results. As the table shows, using default settings for all other parameters, we find that the number of within-species, across-species, and internal HMGTs decreases from 337, 163, and 345, respectively, to 185, 85, and 165, respectively. Nonetheless, even these reduced counts support the widespread presence of HMGTs both within and across species. Furthermore, as can be seen from Supplementary Table S2, all our observations regarding sizes, relative abundances, and patterns of HGTs and HMGTs within- and across-species remain unaffected.

##### Using recipient genome ordering instead of donor genome ordering

By default, we use genome orderings of donor species to infer HMGTs. We repeated the analysis using genome orderings of recipient species instead, and found that nearly all donor-recipient pairs and HMGTs detected using donor species genome orderings are also found when using recipient species genome orderings, and vice versa. For instance, among the 144 within-species donor-recipient pairs inferred using donor genome orderings and 142 inferred using recipient orderings, 136 were in common. Likewise, among the 129 across-species donor-recipient pairs inferred using donor genome orderings and 129 inferred using recipient orderings, 116 were in common. Results are summarised in Supplementary Table S4, which shows that there there is almost no change in the number of donor-recipient pairs and HMGTs detected when using recipient genome orderings.

##### Not skipping over rare genes

In inferring HMGTs using the 〈*x, y, z*〉 parameters, we skip over those genes in the donor genome that occur in small cHGs (or gene families) of size one or two. We refer to such genes as *rare* genes since they are not found in the vast majority of the genomes under consideration, and skip over them because they are likely to have been acquired by HGT from external (or internal) species after an HMGT event. To verify that this choice does not substantially affect HMGT inference results, we performed HMGT analysis without skipping over rage genes. As Supplementary Table S5 show, results remain largely unchanged and we observe a reduction of only 1.2%, 4.9%, and 2.3% in the number of within-species, across-species, and internal HMGTs, respectively.

##### Using specific gene IDs instead of cHGs

Note that each gene present in any of the 103 extant *Aeromonas* genomes has a unique gene ID (or gene name/label), and that each such gene is also associated with exactly one cHG. Thus, each locus in each extant genome has a *gene ID* and a *cHG ID*. In our analysis, we infer HMGTs based on cHG IDs of the transferred genes and the location of genes from those cHGs along the donor (or recipient) genome. This is because specific gene IDs are only available for a subset of detected HGTs (for example, HGTs between ancestral species cannot be assigned to any specific gene ID in extant genomes). As explained in detail in the “Materials and Methods” section, the use of cHG IDs instead of gene IDs, can result in both false-positive and false-negative HMGT inferences. To assess the potential impact of using cHG IDs instead of specific gene IDs, we repeated within-species and across-species HMGT inference using specific gene IDs for donors and recipients of leaf-to-leaf HGTs. Note that it is not always possible to unambiguously infer the specific extant gene ID even for leaf-to-leaf HGTs. On our dataset, out of a total of 39356 within-species HGTs, we were able to infer specific donor and recipient gene IDs for 39041 (or over 99%) of the HGTs. For the vast majority, specifically 38200, of these 39041 HGTs, we could directly determine specific donor and recipient gene IDs because those species each contained only one gene from the corresponding cHG. We were able to infer specific donor and recipient gene IDs for another 841 HGTs by parsing through the gene-tree/species-tree reconciliations used to identify our high-confidence HGTs. For the 14580 total across-species HGTs, we were able to infer specific donor and recipient gene IDs for 13972 (or 95.8%) of the HGTs. As before, gene IDs could be inferred for the vast majority, specifically 12687, of these 13972 HGTs because the donor and recipient species each contained only one gene from the corresponding cHG, and the remaining 1285 HGTs could be assigned specific gene IDs by parsing through the gene-tree/species-tree reconciliations. Thus, we used these slightly smaller sets of HGTs, 39041 within-species and 13972 across species, for the comparative analysis of gene ID based and cHG based HMGT inference.

Supplementary Table S6 shows the results of our analysis. We found that the use of specific gene IDs resulted in slight increases in the numbers of inferred within-species and across-species HMGTs. Specifically, using default values for other parameters, for within-species HMGTs, the number of donor-recipient pairs increased from 143 to 146, with 142 inferred in common, and the number of HMGTs increased from 336 to 344, with 334 in common. Likewise, for across-species HMGTs, the number of donor-recipient pairs increased from 123 to 130, with 121 in common, and the number of HMGTs increased from 157 to 169, with 155 in common. This implies very modest false-positive and false-negative rates of 0.6% and 2.9%, respectively, for within-species HMGTs, and 1.2% and 8.3%, respectively, for across-species HMGTs, when using default parameter settings. Overall, this analysis shows using cHG IDs instead of specific gene IDs has negligible impact on the precision of HMGT inference and minimal impact on recall.

#### Estimating false discovery rate using statistical analysis

If multiple single-gene HGTs have occurred between a donor and recipient, then it is possible for some of those single-gene HGTs to appear next to each other on the donor (or recipient) genome simply by chance. If such a region of contiguous single-gene HGTs is large enough, it may be be falsely inferred to be an HMGT. Such “false” HMGT inferences are more likely to occur as the number of HGTs between a donor-recipient pair increases. We therefore used statistical analysis to estimate the resulting false discovery rate (FDR) of HMGTs and determine appropriate 〈*x, y, z*〉 values to use for our analysis. The analysis is based on randomization of detected HGTs and is described in detail in the “Materials and Methods” section.

Table 2 shows the results of our analysis for four different 〈*x, y, z*〉 parameter choices and reveals several valuable insights. We find that false discovery rates are substantially higher for within-species and internal HMGTs than for across-species HMGTs. This is not surprising since donors and recipients from the same strain have much larger numbers of HGTs (e.g., see Table 1), greatly increasing the likelihood that several of them appear next to each other on the donor and/or recipient genome by chance. We also find that using our default 〈*x, y, z*〉 parameters values of 〈3, 4, 1〉 results in relatively modest FDR estimates, balancing precision with recall. Specifically, we see small FDRs of only 3.95% and 5.34% for across-species pairs and HMGTs, respectively. FDRs are larger for within-species and internal pairs and HMGTs, with 25.83% and 27.48% for within-species pairs and HMGTs, respectively, and 30.14% and 30.38% for internal pairs and HMGTs, respectively. These results also suggest that we likely significantly underestimate the number of HMGTs, particularly across-species and within-species HMGTs, when using our default 〈*x, y, z*〉 parameters values of 〈3, 4, 1〉. For across-species HMGTs in particular, the majority of smaller HMGTs inferred using more permissive parameter values of 〈2, 3, 1〉 are expected to be “true” HMGTs.

**Table 2.** Estimated false discovery rates.

| Within-species |  |  |  |  |  |  |
| --- | --- | --- | --- | --- | --- | --- |
|  | Donor-Recipient Pairs |  |  | HMGTs |  |  |
|  | Rand. Avg. | Actual Total | FDR | Rand. Avg. | Actual Total | FDR |
| X=2, Y=3, Z=1 | 424 | 571 | 74.26% | 1458.6 | 2505 | 58.23% |
| X=3, Y=4, Z=1 | 37.2 | 144 | 25.83% | 92.6 | 337 | 27.48% |
| X=4, Y=5, Z=1 | 4.7 | 40 | 11.75% | 9.5 | 71 | 13.38% |
| X=5, Y=6, Z=1 | 0.4 | 12 | 3.33% | 0.6 | 16 | 3.75% |

| Across-species |  |  |  |  |  |  |
| --- | --- | --- | --- | --- | --- | --- |
|  | Donor-Recipient Pairs |  |  | HMGTs |  |  |
|  | Rand. Avg. | Actual Total | FDR | Rand. Avg. | Actual Total | FDR |
| X=2, Y=3, Z=1 | 66.8 | 551 | 12.12% | 172.7 | 944 | 18.29% |
| X=3, Y=4, Z=1 | 5.1 | 129 | 3.95% | 8.7 | 163 | 5.34% |
| X=4, Y=5, Z=1 | 0.8 | 31 | 2.58% | 0.8 | 36 | 2.22% |
| X=5, Y=6, Z=1 | 0 | 13 | 0.00% | 0 | 14 | 0.00% |

| Internal |  |  |  |  |  |  |
| --- | --- | --- | --- | --- | --- | --- |
|  | Donor-Recipient Pairs |  |  | HMGTs |  |  |
|  | Rand. Avg. | Actual Total | FDR | Rand. Avg. | Actual Total | FDR |
| X=2, Y=3, Z=1 | 842.7 | 966 | 87.24% | 1713 | 2374 | 72.16% |
| X=3, Y=4, Z=1 | 42.5 | 141 | 30.14% | 104.8 | 345 | 30.38% |
| X=4, Y=5, Z=1 | 7.6 | 34 | 22.35% | 15.7 | 76 | 20.66% |
| X=5, Y=6, Z=1 | 1.8 | 9 | 20.00% | 2.4 | 22 | 10.91% |

Note: Results are shown for all $\langle x, y, z \rangle$ parameters settings considered and default settings for all other parameters. The reported randomised average (Rand. Avg.) values for donor-recipient (ordered) pairs and HMGTs are inferred across 10 randomised runs.

For a more fine-grained analysis of FDRs for specific donor-recipient pairs, we repeated the above randomisation analysis separately for each leaf-to-leaf donor-recipient pair. This additional analysis serves to validate that the chosen HMGT inference parameters adequately limit both overall FDR as well as the FDR for any specific donor-recipient pair. We considered all of the potential 571 within-species donor-recipient pairs and 551 across-species donor-recipient pairs (which were identified using the permissive 〈*x, y, z*〉 = 〈2, 3, 1〉 setting; see Table 1) and calculated the fractions of these pairs for which 〈*x, y, z*〉 parameter values of 〈3, 4, 1〉 would yield FDRs of ≤ 5%. We found that our default parameter values of 〈3, 4, 1〉 resulted in an FDR of ≤ 5% for 90.5% (517 out of 571) of the within-species pairs and for 99.5% (548 out of 551) of the across-species pairs. Furthermore, we found that the more permissive 〈*x, y, z*〉 = 〈2, 3, 1〉 setting would have sufficed (for a ≤ 5% FDR) for 73% (402 out of 551) of the across-species pairs, but only for 7.9% (45 out of 571) of the within-species pairs.

#### HMGTs are not functionally biased

We initially hypothesised that genes belonging to certain functional categories would be more likely to be transferred as part of HMGTs, rather than as single genes. To test this hypothesis, we plotted and compared the functional distributions of all transferred genes and genes transferred through HMGTs (i.e., HMGT-genes). Specifically, we assigned each cHG in the analysis to one of 25 COG functional categories (Tatusov et al. 2000), summarised in Supplementary Table S7, and plotted the distribution of these functions separately for all detected HGTs and for all genes transferred as part of inferred HMGTs (inferred using default parameters). Figure 5 shows the results of this analysis for both within-species and across-species donor-recipient pairs. As the figure shows, functional distributions are similar for HGTs and HMGTs, implying that genes from all functional categories are roughly equally likely to be transferred as part of HMGTs. Thus, our dataset and results do not support our initial hypothesis of functional bias. However, we do find that the deviation between functional distributions for HGTs and HMGTs is wider in across-species donor-recipient pairs. In particular, while there is little difference between the within-species plots 5(a) and 5(b), there are some clear differences between across-species plots 5(c) and 5(d) where we find clear cases of over-representation, such as with functional categories [U] and [G], and under-representation, such as with categories [F], [H], [I], and [J], in genes transferred in HMGTs.

**Figure 5.**
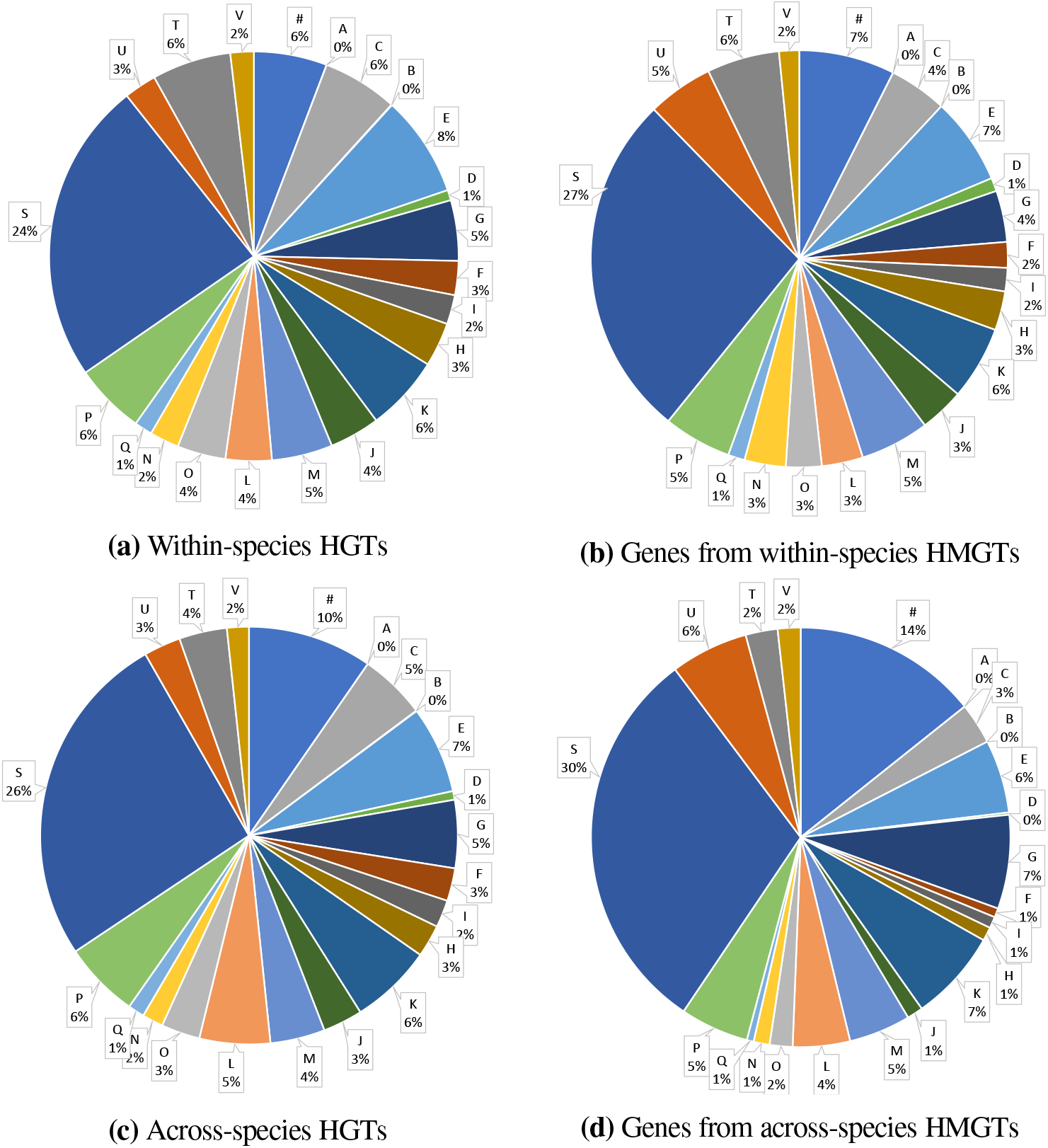
The four pie charts show distributions of COG functional categories for all detected within-species HGTs (part (a)), transferred genes present in within-species HMGTs (part (b)), all detected across-species HGTs (part (c)), and transferred genes present in across-species HMGTs (part (d)). The HGTs and HMGTs used for this analysis were inferred using default parameter settings. Each letter corresponds to a COG functional category, as detailed in Supplementary Table S7. The “#” character labels those genes for which a COG functional category could not be assigned. COG functional categories “Z”, “Y”, “W”, and “R” are not shown in these pie charts since no gene in any of the *Aeromonas* genomes belonged to those categories.

We also plotted the functional distribution of all genes in all 103 genomes and this is shown in Supplementary Figure S5. A comparison of the distributions shown in Figures S5, 5(a), and 5(c) shows that the functional distributions for all detected HGTs, both within- and across-species, are similar to that for all genes in all genomes.

#### Average transfer size increases with increasing phylogenetic distance

Before analyzing this dataset, we had formulated the following hypothesis relating phylogenetic distance and HMGT patterns:

##### Hypothesis 1

*The relative frequency of HMGT, with respect to single-gene HGT, increases with increasing phylogenetic distance*.

In other words, while we expect the absolute numbers of HGTs and HMGTs to decrease with increasing phylogenetic distance (see, e.g., (Williams et al. 2012)), the *relative frequency* of HMGT with respect to single-gene HGT should increase as phylogenetic distance increases. This is consistent with what we observed earlier in Table 1 and Figure 4, where we found that, while the number of HGTs and HMGTs is far higher within-species than across species, across-species donor-recipient pairs have a much higher relative frequency of HMGT than within-species donor-recipient pairs.

To evaluate if our dataset and results support the above hypothesis, we binned inferred donor-recipient pairs by phylogenetic distance and, for each bin, computed the ratio of the total number of genes transferred as part of HMGTs (i.e., the HMGT-genes) and the total number of all detected HGTs for all inferred donor-recipient pairs in that bin. Supplementary Figure S6 shows the results of this analysis. As the figure shows, there is clear trend of increasing relative frequency of HMGTs as phylogenetic distance increases, supporting our hypothesis. For instance, we find that the ratio of HMGT-genes to HGTs is only 0.029 for the first bin (representing the smallest twelfth of phylogenetic distances), averages 0.063 for the two middle bins (bins 6 and 7), and 0.15 for the twelfth bin (representing the largest twelfth of phylogenetic distances). We also repeated this analysis separately for within-species and across-species donor-recipient pairs and results are shown in Supplementary Figures S7 and S8.

Interestingly, we find that the hypothesis holds for across-species donor-recipient pairs but not for within-species donor-recipient pairs. Specifically, we see that relative frequencies of HMGT-genes remain relatively stable across the different phylogenetic distance bins. This is likely due to the fact that within-species phylogenetic distances are very small, with different strains from the same species being nearly identical, providing little meaningful resolution for within-species phylogenetic distance binning.

#### HMGT-genes often have conserved functions

When considering HMGTs, the following question arises naturally: Do HMGTs tend to correspond to meaningful functional units? For example, genes involved in an HMGT may have shared functions or be part of the same pathway. To gain some preliminary insight into this functional aspect of HMGTs, we analyzed inferred within- and across-species HMGTs to determine how often the genes involved in an HMGT were associated with the same COG functional category.

Recall that, with default parameter settings, we infer 337 within-species HMGTs and 163 across-species HMGTs. Among these, we found that 232 within-species HMGTs and 114 across-species HMGTs contained at least one gene with either no function assignment (corresponding to the ‘#’ category in Figure 5) or a “function unknown” assignment of [S]. We therefore limited our initial analysis to just the 105 within-species HMGTs and 49 across-species HMGTs whose genes all had well-defined functions. For the 105 within-species HMGTs, we found that 48 (45.7%) of the HMGTs had distinct functional assignments for each of their genes (i.e., in the detected transferred genes present in those HMGTs), 47 (44.76%) of the HMGTs has the same functional assignment for more than half of their genes, and 21 (20%) of the HMGTs had the same functional assignment for all their HMGT-genes. Interestingly, across-species HMGTs showed much greater functional conservation. Specifically, among the 49 across-species HMGTs, only 8 (16.3%) had distinct functional assignments for each of their HMGT-genes, 36 (73.47%) had the same functional assignment for more than half of their HMGT-genes, and 18 (36.73%) had the same functional assignment for all their HMGT-genes.

Supplementary Figure S9 plots some of these results and highlights the considerable difference between functional conservation patterns in within-species HMGTs and across-species HMGTs. This difference may reflect the mode by which genes are integrated into the recipient genome: Homologous recombination, which is the dominant integration mechanism expected for within-species HGTs/HMGTs, does not limit the transferred genes to functional units beyond the neighborhood relations in the genomes; in contrast, genes transferred across species often are selfish genetic elements, pro-phage, and genomic islands, whose individual genes often fall into the same functional category. Another factor may be the detectability of within-species HGTs/HMGTs. To detect HGTs using phylogenetic conflict, the sequences need to have accumulated polymorphisms. Well-characterized genes are often under stronger purifying selection than genes without assigned function. As a consequence, the within-species transfer of a group of genes under strong purifying selection, such as ribosomal proteins or ATP synthase subunits, will not be detected using phylogenetic conflict as a criterion.

We note that this difference in functional conservation patterns persists even if all HMGTs are considered, treating ‘#’ and [S] as “functions”. Specifically, among all 337 within-species HMGTs, the numbers of HMGTs with no functional conservation, >50% functional conservation, and 100% functional conservation are 154 (45.7%), 140 (41.5%), and 36 (10.7%), respectively. For all 163 across-species HMGTs, the corresponding numbers are 37 (22.7%), 103 (63.2%), and 35 (21.5%), respectively.

To further ascertain the significance of the functional conservation trends noted above, we performed statistical analysis to determine if the genes present in an HMGT were assigned the same COG functional category more often than would be expected by chance. For this analysis, we randomised the functions of the genes in the inferred within-species and across-species HMGTs and calculated, as above, the number of HMGTs with >50% functional conservation and 100% functional conservation. The randomised functions were drawn from the overall functional distribution of detected HGTs (after removing genes assigned ‘#’ and [S] categories), separately for within-species HGTs and across-species HGTs, and the randomised analysis was repeated 100 times. For the 105 within-species HMGTs, the randomisation analysis resulted in an average of 16.4% and 0%, respectively, of HMGTs with >50% functional conservation and 100% functional conservation. Even the maximum counts of within-species HMGTs with > 50% functional conservation and 100% functional conservation, observed among the 100 randomised runs, were only 26.7% and 0%, respectively. Likewise, for across-species HMGTs, the randomisation analysis resulted in an average of 18.4% and 0%, respectively, of HMGTs with >50% functional conservation and 100% functional conservation. The maximum counts of across-species HMGTs with >50% functional conservation and 100% functional conservation, observed among the 100 randomised runs, were only 32.6% and 0%, respectively. This statistical analysis shows that the observed levels of functional conservation in within- and across-species HMGTs are highly unlikely to have occurred by chance (*p* < 0.01).

#### Qualitative Analysis of HMGTs

Although the quantity and functional ratios differed, all of the functional groups discussed here are present in both across and within species inferred transfers. An examination of these lists reveals that many of the genes transferred are those known to be commonly transferred (Nakamura et al. 2004, Zhaxybayeva et al. 2006). For instance, there were large numbers of phage related genes including major capsid proteins, tape measure proteins, terminases, and phage integrases to name but a few. These phage genes were often flanked by additional genes of no known function or occasional virulence factors (e.g., Zonula Occludens Toxin). There were also several kinds of transposable elements transferred within our dataset including the Tn7 and several unclassified transposition proteins. Bacterial defense mechanisms were also commonly transferred genes. Among this group, the least common were anti-phage systems. These included: one CRISPR cassette, three sets of restriction modification system genes (all type I), and a number of restriction endonucleases. Much more prominent were the transfers of antimicrobial resistance genes. Transfers involving these resistance genes often included transposition proteins as part of the HMGT, which suggests transposons as the main means of transfer. These resistance genes included beta-lactamases, tetracycline resistance genes, achloramphenicol acetyltransferases, tetracycline resistance proteins, and a polymixin resistance gene. Finally there were several transfers of virulence genes (e.g., type III secretion system) numerous transfers of metabolic enzymes (e.g., nudix hydrolase, pseudouridine synthase), and many transporters (principally ABC transporters).

#### HMGT of Zonula Occludens Toxin genes

Of particular interest to us were HGTs of virulence related genes. One of these genes, the Zonula Occludens Toxin (ZOT), caught our early attention. The toxin is well known for its role within *Vibrio cholerae*, where it acts to disrupt intacellular signalling and break up tight junctions (Pierro et al. 2001). It is also known to as part of the CTXΦ phage (Baudry et al. 1992, Waldor and Mekalanos 1996) which helps to transfer the toxin between various *Vibrio* strains (Boyd et al. 2000). Our initial results indicated that the ZOT from cHG 11010 was being transferred in an HMGT with two other genes which we will refer to as 1729 and 1929. Extensive database searches and our RAST annotation results indicated that the genes 1729 and 1929 coded for a viral period protein, and a minor coat protein (specifically a VSK receptor), respectively.

Investigation of the syntenic regions surrounding this inferred transfer garnered two crucial observations. First being that there were three different ZOTs in three separate cHGs present in our genomes. A phylogeny of the three cHGs (11010, 14858, and 4422) with toxins samples from outside the *Aeromonas* revealed that each was a divergent copy of the same toxin (Supplementary Figure S10). Outgroup accession numbers are available in Supplementary Table S12. Analysis of the syntenic regions showed that all three groups integrated at the same syntenic region in the genome. Specifically all of the toxins could be found between the YebG SOS response gene and a 3-hydroxyacyl dehydratase (3HD), except in two cases where the synteny of the region was disrupted, likely as a result of homologous recombination. In a few instances this site was home to multiple copies of the toxin across the three cHGs (for example, *Aeromonas veronii* BAQ135 had a copy of the toxin from each of the three cHGs). Investigation of the region between these YebG and 3HD uncovered the consistent inclusion of phage integrases and replication initiation factors adjacent to the YebG and 3HD respectively. Between the integrase and replication initiation factor were various hypothetical and known viral proteins; however, as Figure 6 shows, the ordering was rarely conserved between genomes. This indicates that this toxin is likely moving through *Aeromonas* with the assistance of a phage similar to CTXΦ over periods of time long enough to allow for the recombination of regions within these prophages. A previous study found similar transfers of a large element containing the toxin (Tekedar et al. 2019) which provides further support for this hypothesis.

**Figure 6.**
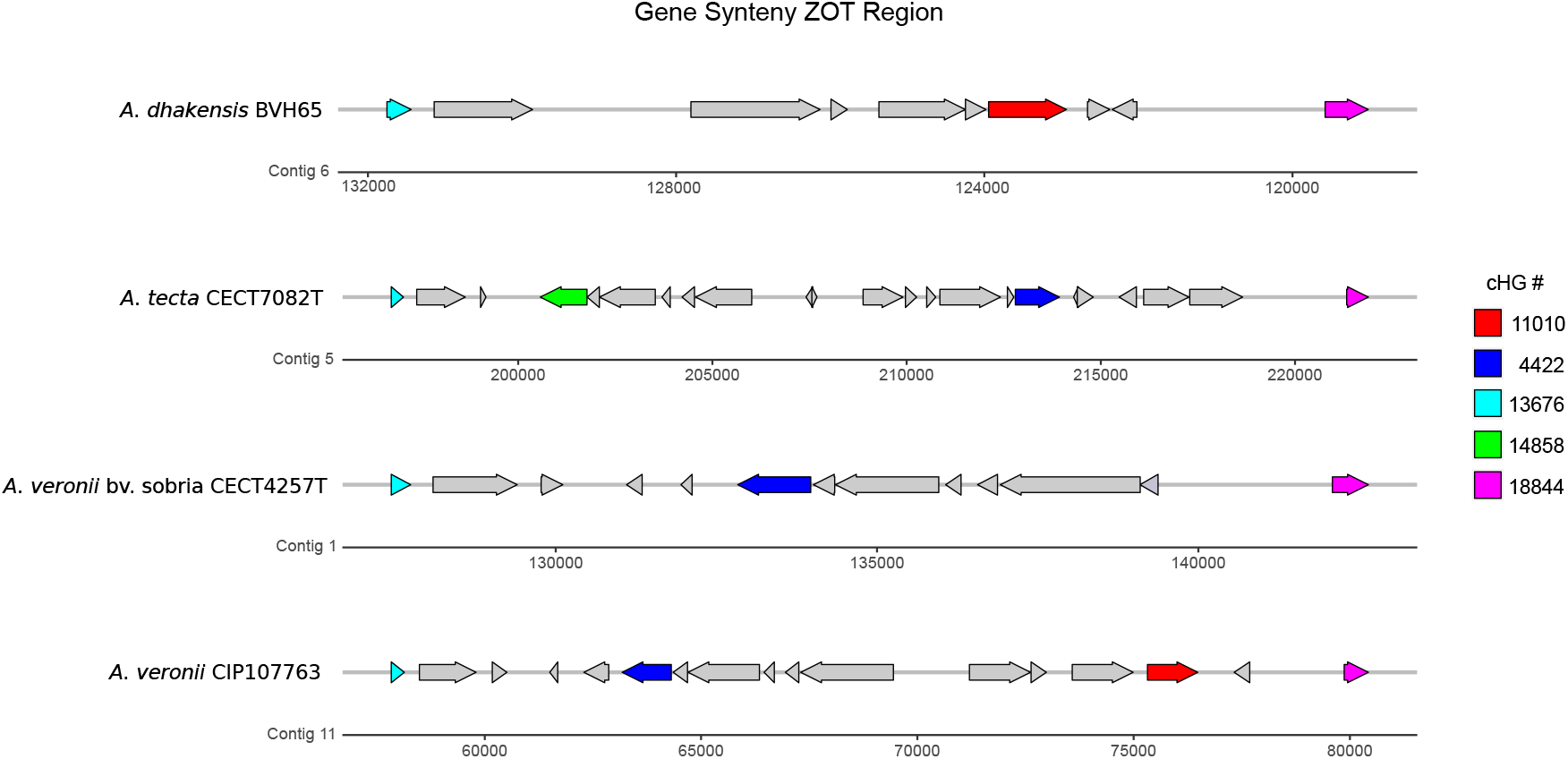
Gene synteny plot depicting the diversity of genes and their synteny within the ZOT integration site. Colored in cyan is YebG (cHG 13676) and in magenta is 3HD (cHG 18844). All other colored cHGs are different versions of the ZOT gene. Arrows depict coding direction. The given numbers correspond to contigs in the draft genomes. For information on all genes present within this plot see Supplemental Figure S11.

#### HMGT of type III secretion systems

We also used this dataset as an opportunity to expand upon prior work on horizontal transfer of the type III secretion system (T3SS) within the *Aeromonas* (Rangel et al. 2019). The T3SS acts as a molecular syringe which acts to transfer effector proteins (with a wide range of possible biochemical activities) into cells (Dean, 2011). The genes associated with this system have been shown to frequently horizontally transfer and are often found within pathogenicity islands (Hacker et al. 1997). Within the *Aeromonas*, there are two different forms of the T3SS that have been found previously to be transferred around the genus (Rangel et al. 2019); we refer to these as T3SS-1 and T3SS-2.

We examined the list of across-species HMGTs (inferred using default parameters) for any instances of T3SS within the annotations. We found there were 7 HMGTs which contained genes pertinent to the T3SS. These HMGTs are shown in the colored boxes in Supplementary Figure S12. Of these 7 HMGTs, 3 pertain to T3SS-1 and 4 pertain to T3SS-2. In a similar fashion to the ZOT, the syntenic regions around these cHGs were investigated for any consistent marker genes; however, a visual search did not unveil any obvious and consistently occurring flanking genes. As for the syntenic region itself, the sites around these inferred HMGTs rarely had similar syntenic compositions. The inferred HMGT-genes were then used as the base for creating gene trees such that we could investigate the relationship of *Aeromonas’s* T3SS to other closely related species. Each cHG served as the base for its respective gene tree, constructed using NCBI’s non-redundant database, MUSCLE, and IQTree (see “Materials and Methods” Section for additional detail). These gene trees are shown in Supplementary Figure S13. Next, we describe our major findings from this analysis.

In the T3SS-1 phylogenies, most *Aeromonas* cluster closely together with very high bootstrap support; however, *A. schubertii* consistently groups sister to the *Yersinia* genus in every cHG. Furthermore, in some gene trees there are other *Aeromonas* which group with *schubertii* and sister to the rest of *Yersinia* (for example, in gene tree 9090 A. *tecta* CECT7082T and *A. veronii* AMC25 both group with *schubertii* sister to *Yersinia*). This may indicate that *schubertii* has, after acquiring its T3SS from *Yersinia*, transferred genes from this more divergent T3SS-1 into other members of the genus. Otherwise it is possible that these cases were separate HGT events from *Yersinia* species to other *Aeromonas* taxa.

In the T3SS-2 phylogenies, the few *Aeromonas* present cluster closely with *Salmonella enterica* subspecies enterica. With one major exception, the T3SS-2 in *Aeromonas* appears to be a more recent acquisition among those that possess it, as there is very little to no variation in their sequences despite the distance separating them on our species tree. The exception to this is *A. jandaei* Ho603. This taxon is on a consistently long branch, and more often than not separated from the rest of the *Aeromonas* by several interior nodes. It is not clear where this divergent version of the T3SS-2 comes from, as there is no consistent grouping for this *jandeai* strain. However, its divergence is such that manual blast searches (with less stringent e-value cutoffs and smaller gap penalties) were necessary to find matches for cHGs 369, 803, 11915 and 19118 (and the two that had matches were once again extremely divergent copies).

#### Rare genes are frequently acquired through HMGT

Note that genes from small cHGs that have only one or two genes are not included in the results described above. Such genes, which we refer to as *rare genes*, were likely acquired through HGT from species not represented in the 103-genome *Aeromonas* species tree. There are a total of 15965 rare genes from 13524 cHGs in our dataset. To determine if any of these rare genes may have been acquired via multi-gene transfer, we mapped the location of each rare gene on each extant genome ordering and, for each extant genome, inferred putative HMGTs comprising of rare genes using various 〈*x, y, z*〉 parameter values. Observe that the donor species for rare gene transfers are assumed to be unknown, and the extant genomes serve as recipient species for this HMGT analysis.

Supplementary Figure S14 shows the distribution of rare genes in the 103 genomes. On average, the genomes contained 155 rare genes, with a maximum of 669. Since many of the genomes have high numbers of rare genes, resulting in high FDR, we report here results of analyzing just the 40 genomes that each contained less than 100 rare genes. For completeness, results for all 103 genomes are available in Supplementary Table S8.

Table 3 shows the results of our analysis. We find that a large fraction of the rare genes were likely acquired together with other rare genes through HMGTs. For instance, with our default 〈*x, y, z*〉 parameter setting of 〈3, 4, 1〉, 34 out of the 40 genomes were found to have rare-gene HMGTs, with a total of 107 rare-gene HMGTs distributed across those 34 genomes. These HMGTs contained a total of 469 rare genes, representing 19.7% of all rare genes present in these 37 species. These results are consistent with our previous results on across-species HMGTs (see Table 1), where we also found roughly 20% of detected HGTs being transferred as part of larger HMGTs. Notably, we also found that many of the detected rare-gene HMGTs were quite large, with the 15 largest rare-gene HMGTs containing an average of almost 9 rare genes each.

**Table 3.** Results of rare-gene HMGT inference analysis for the restricted set of 40 *Aeromonas* genomes.

| Parameters | Genomes<sup>1</sup> | Rare-Gene HMGTs<sup>2</sup> | HMGT-genes<sup>3</sup> | Total Rare Genes<sup>4</sup> |
| --- | --- | --- | --- | --- |
| $x=2, y=3, z=1$ | 40 | 365 | 991 | 2566 |
| $x=3, y=4, z=1$ | 34 | 107 | 469 | 2371 |
| $x=4, y=5, z=1$ | 24 | 54 | 305 | 1835 |
| $x=5, y=6, z=1$ | 18 | 31 | 213 | 1474 |
| $x=6, y=7, z=1$ | 9 | 15 | 131 | 765 |

Note: Results are shown for all $\langle x, y, z \rangle$ parameters settings considered and default settings for all other parameters.
<sup>1</sup> The number of genomes that had at least one rare-gene HMGT.
<sup>2</sup> Total number of inferred rare-gene HMGTs.
<sup>3</sup> Total number of rare genes present within the inferred rare-gene HMGTs.
<sup>4</sup> Total number of rare genes present in the corresponding genomes.

We also performed statistical analysis to estimate FDRs for this rare-gene HMGT analysis. This statistical analysis was performed along similar lines as before, with genes selected randomly from each genome. Specifically, for each of the 40 genomes, we randomly sampled as many genes as the number of rare genes in that genome and applied our HMGT inference pipeline using these randomly chosen genes. This analysis supports our results, showing that the expected FDR for rare-gene HMGTs using our default 〈*x, y, z*〉 parameter setting of 〈3, 4, 1〉 is only 1.5%. Even with the more permissive parameter setting of (2,3,1), the FDR is only 23.3%. Complete results of the statistical analysis appear in Supplementary Table S10.

We point out that results on the complete set of 103 genomes are consistent with the results reported above for the 40-genome analysis. Specifically, we see 778 rare-gene HMGTs containing a total of 3870 rare genes across 97 of the 103 genomes when using default parameter settings of (3,4,1) (Supplementary Table S8), with an estimated FDR of 18.25% (Supplementary Table S9. Consistent with what we observed above with the 40-genome analysis, these 3870 rare genes contained within rare-gene HMGTs represent 24.5% of all rare genes present in the 97 species (Supplementary Table S8).

#### Rare-gene HMGTs show interesting functional characteristics

To determine if rare genes and rare-gene HMGTs had different functional distributions than for regular HGTs and HMGTs, we analysed the COG functions for rare genes and rare-gene HMGTs from the 40 chosen genomes with less than 100 rare genes. Strikingly, we find that most rare genes could not be matched to any COG functional category. This is depicted in Supplementary Figure S15(a), which shows that 55% of all rare-genes could not be assigned a COG category. This observation holds even for the genes within rare-gene HMGTs, with 52% not assigned to any COG functional category (Supplementary Figure S15(b)). In contrast, only 6% of HGTs and 7% of HMGT-genes had no assigned COG functional category. This great over-representation of genes with no matching COG category among the rare genes has at least two possible explanations: (a) these genes have a low frequency of occurrence, not only in Aeromonads, and therefore these genes and their homologs may not have been characterized to date; (b) some of these genes might be gene calling artifacts. The latter is less likely for HMGTs, since gene calling mistakes for several sequential genes are less likely than for a single gene.

Comparing functions of rare-genes with functions of genes within rare-gene HMGTs, we find that there is clear over-representation of genes from the [L] functional category (replication, recombination and repair) among rare-gene HMGTs. As Supplementary Figure S15(b) shows, 10% of the genes present in rare-gene HMGTs belong to the [L] functional category, while only 5% of all rare genes belong to that category. This is not surprising since we see many genes of phage and selfish genetic elements in rare HMGTs, and transposases, integrases, components of restriction modification systems, conjugative transfer proteins are all placed into the [L] category.

Some other interesting observations related to rare-gene HMGTs include: (i) an abundance of glycosyl transferases and other enzymes likely involved in modifying the bacterial surface (e.g., colanic acid biosynthesis, rhamnosyltransferase) and in carbohydrate metabolism (e.g., maltooligosyl trehalose trehalohydrolase), (ii) presence of genes that appear to encode enzymes in the synthesis and modification of secondary metabolites (e.g., nikkomycin biosynthesis, biosynthesis of phenazines, glyoxalase/bleomycin resistance protein/dioxygenase), and (iii) a cluster of two likely heme agglutinine genes in *A.hydrophila-CECT839T*.

## Discussion

In this work, we introduced a new computational framework, HoMer, for the systematic discovery of HMGTs at a large scale. Its application to the Aeromonads demonstrates the prevalence of HMGTs as well as their significance to microbial evolution. For instance, we found that HMGTs are ubiquitous and a large fraction of transferred genes are transferred as part of HMGTs, at both short and large phylogenetic distances. We also found that the relative frequency of HMGT increases as divergence between genomes increases, that HMGTs often have conserved gene functions, that genes from all functional categories appear to be roughly equally likely to be transferred as part of HMGTs, and that rare genes acquired from outside a particular clade of interest are frequently acquired through HMGT. Our analysis of HMGTs involving the ZOT and T3SS shows that within-genus HMGTs play an important role in diversifying host-symbiont interactions, and that in the case of the ZOT, phages appear to play a major role in shuffling the ZOT gene neighborhood via repeated recombination and invasion events.

HoMer is easy to use, scalable, and effective, and makes it feasible to systematically infer HMGTs on a large scale. We expect that the systematic discovery of HMGTs, enabled by this work, will lead to enhanced understanding of horizontal gene transfer and microbial evolution. Nonetheless, the current HMGT inference framework implemented in HoMer has some limitations and potential biases worth understanding. A key limitation is that our ability to infer HMGTs depends on there being sufficient phylogenetic resolution in the gene trees to reasonably detect (single-gene) HGT events. This limitation makes it harder to infer HGTs and HMGTs between closely related pairs of strains or species, and can thus bias HMGT inference results by resulting in a greater false negative rate for such pairs. Another important limitation is that our approach is focused on finding HMGTs that are “large enough” to be unlikely to occur by chance. In other words, to control for the false discovery rate, the 〈*x, y, z*〉 parameter values have to be set conservatively. However, as our statistical analysis (Table 2) suggests, the vast majority true HMGTs may be smaller than are detectable using our default 〈*x, y, z*〉 parameter setting of 〈3, 4, 1〉.

While our experimental analysis with the Aeromonads sheds light on the prevalence of HMGT and provides several fundamental insights, many important questions remain unanswered. For instance, it would be interesting to investigate if HMGTs tend to correspond to operon boundaries or to functional pathways. It would also be useful to extend our computational framework to make it more suitable for detecting HMGTs between more distantly related species with little gene order conservation. This may be achieved by combining HoMer with methods that model genome rearrangement and/or infer ancestral genome orderings.

## Materials and Methods

### Dataset construction

103 previously published complete and draft *Aeromonas* genomes were used in this study (Rangel et al. 2019). Protein coding ORFs were called by the RAST annotation server (Aziz et al. 2008). Genome completeness, GC content, size, and other statistics were calculated using CheckM (v1.0.7) via the taxonomy_wf option and the Aeromonadaceae as the family database (Parks et al. 2015). Supplementary Table S14 shows a complete listing of genomes along with related statistics. As the table shows, these 103 genomes had an average completeness score of 99.47%, and only one genome had a completeness score below 97.9%. These genomes had a average of 93.5 contigs, with a median value of 67, and only 23 genomes had more than 100 contigs.

The reference species tree was inferred via the 16-gene multi-locus sequence analysis (MLSA) scheme previously established for use in the *Aeromonas* (Colston et al. 2014). Sixteen single-copy housekeeping genes were extracted via BLAST and concatenated into a single alignment as described in Colston et al. (2014). The phylogeny was inferred using RAxML (v. 8.1.21) under a GTR+GAMMA+I model (Stamatakis 2014). Consistent with previous analysis of the Aeromonads (Colston et al. 2014, Rangel et al. 2019), we rooted the species phylogeny along the branch connecting the *A. schubertii, A. diversa, A. simiae* clade to the rest of the tree.

Details on homology clustering, generation of cHGs, functional annotation, synteny mapping, and data related to the ZOT and T3SS analyses appear in the Supplement (Section titled “Supplementary Text: Dataset Construction”).

#### Basic statistics on dataset

The 103 genomes in the dataset correspond to 28 different species. Of these 28 different species, 18 are represented by a single strain (genome), while the remaining 10 are each represented by at least two genomes corresponding to different strains from that species. Supplementary Figure S1 shows the distinct species that appear in the dataset along with the number of genomes/strains representing each species. The full species tree topology is shown in Supplementary Figure S2.

Of the total of 22282 cHGs, 8277 had at least three genes and the remaining 14005 cHGs had either a single gene or two genes. We were thus able to construct gene trees for 8277 cHGs. The average size of these 8277 cHGs was 48.8 genes. The average number of genes per genome was ~ 4090, of which roughly 96%, on average, were represented in one of the 8277 gene trees. The remaining ~ 4% of genes, corresponding to cHGs of size 1 and 2, were aggregated into a list of “rare” genes and analyzed separately as described in the “Results” section.

### Methodological details

We describe the key steps of HoMer in detail below.

#### Inference of high-confidence HGTs

We used phylogenetic reconciliation of gene trees with the species trees to infer HGTs on the species tree. To construct the gene trees used for reconciliation, protein sequences were backtracked to DNA sequences via Perl scripting and the RAST-generated genomic spreadsheet files, and sequences within each cHG were aligned with MUSCLE (v3.8.31) (Edgar 2004). Gene trees were constructed on these aligned sequences using RAxML (Stamatakis 2014) (GTR+GAMMA+I model, thorough search settings with 100 rapid bootstraps per tree) and these RAxML trees were further error-corrected using TreeFix-DTL (Bansal et al. 2015) (GTR+GAMMA+I model, default search settings). TreeFix-DTL essentially removes statistically unsupported differences between the gene tree and species tree, making the final set of inferred HGTs more accurate (Bansal et al. 2015).

Phylogenetic reconciliation was performed using RANGER-DTL 2.0 (Bansal et al. 2018) which reconciles gene trees with species trees by invoking gene duplication, gene loss, and HGT events. We used RANGER-DTL 2.0 (Bansal et al. 2018) to optimally root the TreeFix-DTL gene trees and compute optimal DTL reconciliations. To account for reconciliation uncertainty, we sampled 100 optimal reconciliations per gene tree and aggregated across these samples to identify highly-supported HGT events. The specific support thresholds used in our analysis are reported in the next subsection.

Note that only cHGs with at least three genes were used for reconciliation based HGT inference. Those cHGs with only one or two genes were analyzed separately (see “Results” Section).

#### Mapping HGTs to genomic locations

Each high-confidence HGT event inferred through the steps above is associated with a specific donor and recipient species on the species tree. For each possible donor-recipient pair in the species tree, we (i) assemble a list of all cHGs that have an HGT from that donor to that recipient and (ii) mark the locations of those transferred genes along the donor (and/or recipient) genome(s) using the available gene ordering information. Since gene orders are only available for extant species, we perform step (ii) only for HGTs where the donor and recipient are both leaves (i.e., extant species) on the species tree.

#### Defining HMGTs for transfers between extant species

We define HMGTs to be regions of the donor and/or recipient genome that have “unusually many” high-confidence HGTs clustered together. Identification of HMGTs is complicated by the fact that any collection of inferred HGTs is expected to have relatively high false-positive and false-negative rates. For instance in our analysis we focus on using only “high-confidence” HGTs and therefore expect a relatively high false-negative rate. Moreover, there can be considerable uncertainly in correctly identifying the donor and recipient species for individual HGT events. We therefore define HMGTs formally using three parameters 〈*x, y, z*〉, where we first identify contiguous regions of *y* genes in which at least *x* genes were detected as transferred from the donor to the recipient, and then merge the identified regions with neighboring regions or HGTs if the distance between them is no more than z. For appropriately chosen values of 〈*x, y, z*〉, e.g., 〈3, 4, 1〉, each of these merged regions constitutes a plausible HMGT. This is illustrated in Figure 1.

In defining these plausible HMGTs, we also account for the presence of rare genes that occur very infrequently in the genomes of the considered set of species. More precisely, when identifying plausible HMGTs using the 〈*x, y, z*〉 parameters, we skip over all those genes in the donor (and/or recipient) genome that occur in cHGs of size one or two. This is based on the observation that such rare genes may have been acquired by HGT from external (or internal) species after an HMGT event and, consequently, should not be allowed to disrupt the detection of those HMGT events. As described in the “Results” section, skipping over such rare genes has a small, but non-negligible, impact on HMGT inference.

Details on the statistical analysis used to select appropriate 〈*x, y, z*〉 parameters and estimate the false positive rate for inferred HMGTs appear below in the subsection titled “Statistical analysis”.

#### Defining HMGTs for transfers that are not between extant species

Since gene orders are only available for extant species, to infer HMGTs when at least one of the donor or recipient is an ancestral species, we look for plausible HMGTs using the most compliant ordering of any of the extant descendants of the donor species (or recipient species). Specifically, for the inferred set of transfers, we compute the number of HMGTs implied by each of the leaf descendants of the donor species (or recipient species). The leaf descendant that implies the largest number of HMGTs is then used for inferring all HMGTs for that donor (or recipient). Essentially, the goal is to identify and use that leaf descendant whose gene ordering is likely most similar to that of the actual donor (or recipient) species.

### Specific parameter choices

HoMer provides many parameters that can be fine tuned to adjust the sensitivity and specificity of the analysis and control for the kinds of HMGTs that are discovered. These parameters can be broadly divided into those related to HGT inference and those related to HMGT inference. We describe and justify below the specific parameter settings used in our analysis of the *Aeromonas* dataset.

#### HGT inference parameters

We used stringent parameter choices for HGT inference so as to obtain conservative estimates of the prevalence of HMGTs. For our primary analysis, we used duplication, transfer, and loss costs of 2, 4, and 1, respectively. Transfers are typically assigned a cost of 3 when performing DTL reconciliation (David and Alm 2011, Bansal et al. 2012; 2013a), and a higher cost of 4 implies that HGTs are only invoked where an alternative scenario involving duplications and losses is unlikely. The resulting increase in precision comes at the cost of a slight decrease in specificity since HGTs between very closely related species may be missed. For comparison, we also report results using the default duplication, transfer, and loss costs of 2, 3, and 1, respectively (Bansal et al. 2018).

To account for reconciliation uncertainty and identify unambiguous HGTs, we sampled 100 randomly chosen optimal reconciliations per gene tree and aggregated across these samples. We only chose those HGTs that had 100% support (i.e., HGTs that were present in all 100 sampled reconciliations for that gene tree). Even for an unambiguous HGT there is often uncertainty about its exact donor and recipient species (i.e., originating and receiving edge on the species tree) (Bansal et al. 2013a). To account for such uncertainty, we further filtered the set of unambiguous HGTs to those that had a consistent donor assignment (mapping) and a consistent recipient assignment (mapping) across at least 51% of the sampled optimal reconciliations each. This 51% threshold was chosen so as to balance the demands of identifying a well-supported donor and recipient for each unambiguous HGT, while not discarding too many unambiguous HGTs. To assess the impact of this parameter choice, we also computed results with a 75% donor and recipient mapping threshold.

These parameter choices are summarized in Supplementary Table S1.

#### HMGT inference parameters

The most impactful parameters for HMGT inference are the 〈*x, y, z*〉 parameters used to define what constitutes an HMGT. We used a default setting of 〈3,4,1〉 for these parameters. With this setting, each identified HMGTs must have involved the simultaneous transfer of at least three syntenic genes, and often of four or more syntenic genes. This default setting was chosen because it results in a relatively small false discovery rate (see section on “Statistical analysis” below for details on how false discovery rates were estimated). To estimate the number of larger HMGTs as well as putative smaller HMGTs we also computed HMGTs with parameter values 〈2,3,1〉, 〈4,5,1〉 and 〈5,6,1〉, as well as with *z* increased to 2.

As described above, to mitigate any confounding effects of recently transferred rare genes we chose to skip over rare genes when detecting HMGTs using genome orderings. For comparison we also computed results without skipping over such genes.

For any given donor-recipient species pair, the genome ordering of either the donor species or the recipient species can be used to detect plausible HMGT events. By default, we chose to use donors’ genome orderings to compute HMGTs. To assess the impact of this choice on HMGT detection, we also repeated our analysis with recipients’ genome orderings.

For each species, genome orderings are available as ordered lists, one per contig, of gene IDs from that species. For each gene ID in such an ordering, the cHG it belongs to is also known. Note that only the genes in extant genomes have specific gene IDs or labels. Consequently, the specific gene transferred in an HGT event is generally not known (e.g., an HGT may transfer some ancestral gene from some ancestral species). As a result, in general, for any HGT event, we only know its donor, recipient, and the cHG the transferred gene belonged to. Thus, in using the extant genome orderings to detect HMGTs, by default, we view the genome orderings as ordered lists of gene families rather than as ordered lists of specific gene IDs. This can be slightly problematic in certain cases since genomes may have multiple genes from the same cHG, implying that the same cHG may occur multiple times in a single genome ordering. In our implementation, we only consider the last occurrence of a cHG in a genome ordering and ignore any previous occurrences. This can give false negatives when detecting HMGTs, since we may not be looking at the “correct” location of that cHG in the genome ordering. In very rare cases, this can also lead to false positives, when an “incorrect” location of that cHG nonetheless yields a putative HMGT. To estimate the number of false negatives and false positives resulting from the use of cHG IDs rather than specific gene IDs, we also used a subset of recent HGTs for which specific extant gene IDs could be inferred and computed HMGTs based on those gene IDs.

These parameter choices are summarized in Table S1.

### Statistical analysis

Given any donor-recipient pair, it is possible for several single-gene HGTs to appear clustered together on the donor (or recipient) genome by chance. Such “false” HMGTs are more likely to occur as the number of HGTs for that donor-recipient pair increases. We therefore used statistical analysis to estimate the resulting false discovery rate (FDR) of HMGTs and determine appropriate 〈*x, y, z*〉 values to use for our analysis.

For any fixed setting of HGT and HMGT inference parameters, we estimate the resulting FDR by first randomizing the inferred HGTs while preserving total HGT counts as well as their donors and recipients, and then executing the same HMGT inference pipeline using these randomized HGTs instead of inferred HGTs. Specifically, for each pair of donor-recipient edges (*e_i_, e_j_*) on the species tree, let denote the set of HGT events inferred with *e_i_* as donor and *e_j_* as recipient. Let *G_ij_* denote the set of cHGs shared between the species represented by edges *e_i_* and *e_j_*. Then, for each such (*e_i_, e_j_*), we randomly choose the same number |*H_ij_*| of HGTs from the collection of *G_ij_* shared cHGs. We then infer HMGTs using these randomized HGTs and record how many HMGTs were inferred. This gives an estimate of the FDR of HMGTs for that specific setting of HGT and HMGT inference parameters. For improved accuracy, we repeat this randomization analysis 10 times, for each setting, and average over the results.

Observe that the above randomization analysis yields an estimate of the FDR for HMGTs over the entire species tree. For specific donor-recipient pairs, the expected FDR could be smaller or larger than this overall FDR. We therefore also repeated the above analysis separately for each donor-recipient pair. This additional analysis serves to validate that the chosen HMGT inference parameters don’t just adequately limit overall FDR but also sufficiently control FDR for any specific donor-recipient pair. This analysis also helps identify those donor-recipient pairs for which the overall HMGT inference parameters could be made more permissive, and also to identify donor-recipient pairs for which the chosen HMGT inference parameters may be too permissive.

## Data availability

The genomic data (i.e., complete and draft *Aeromonas* genomes) underlying this article are all publicly available (Rangel et al. 2019). The gene families, gene trees, gene ordering information, species tree, and software used for our analysis are freely available from https://compbio.engr.uconn.edu/software/homer/.

## Acknowledgments

This work was supported by the US National Science Foundation (grant number MCB 1616514 to M.S.B., J.P.G., and J.G.) and United States Department of Agriculture (CRIS project award 8082-32000-006-00-D to J.G. and J.P.G.).

## Notes

https://compbio.engr.uconn.edu/software/homer/

### Competing Interest Statement

The authors have declared no competing interest.

