## Supplementary text, tables, and figures for "Systematic Detection of Large-Scale Multi-Gene Horizontal Transfer in Prokaryotes"

#### Supplementary Text: Dataset Construction

**Homology clustering.** Protein sequences from the 103 *Aeromonas* genomes, plus one later discarded, were clustered into homologous groups using OrthoMCL (Li et al. 2003) as implemented in the GET_HOMOLOGUES software package (Contreras-Moreira and Vinuesa 2013). A total of 25518 homologous groups were assembled, 2755 of which are present in at least 90% of surveyed genomes. The genome of *A. enteropelogenes* 1999lcr was then discarded on account of displaying both an excessive number of contigs (1139) and low N50 value (5421). It likely suffered from an elevated number of annotation artefacts and would have reduced the chance of extracting meaningful syntenic information. Removal of its member ORFs without a fresh *de novo* clustering was deemed acceptable as these initial clusters would be clustered into larger consolidated groups in the next step of analysis.

**Collapsing of clusters to generate cHGs (or gene families).** It has been observed that GET_HOMOLOGUES is more likely to over-split groups under default parameters than to under-split them. As the methodologies described below are adept at coping with paralogy and large datasets, many of the clusters were collapsed. The longest ORF in each cluster was extracted for a secondary search to stand as a proxy for the entire HG. The ublast function of the USearch (v. 8.0.1623) was used with identity of 0.3, target and query coverages of 0.6, and an e-value cutoff of 1e-8 to search each longest ORF against each other longest ORF (Edgar 2010). The resulting hits were loaded into the igraph package (Csardi and Nepusz 2006) in R. The components function effectively generated a single-linkage clustering of the HGs, grouping HGs meeting the ublast criteria into larger clusters. The exported list was then fed into Perl scripting that combined the HGs into consolidated HGs (cHGs). This resulted in a final total of 22282 cHGs, also referred to as gene families.

**Functional assignments for cHGs.** A representative member was randomly chosen from each cHG and copied over to create a multifasta file containing representatives from all cHGs. These nucleotide sequences were translated into amino acid sequences. The hmmscan function of HMMER (v3.2.1) (`http://hmmer.org`) was used to query these sequences against the Trinotate customized version of the Pfam database (Bryant et al. 2017). Final annotations were extracted from HMMER outputs via in house Perl scripting.

**Synteny Mapping.** In-house Perl scripting was used to extract meaningful synteny information from RAST output files. This data was organized such that: each synteny file covers a single genome, each line covers one contig, each ORF is tab delimited, and within each ORF data field is listed: peg/ORF ID, contig ID, strand (+/-), RAST functional calls, and consolidated homologous group number to which the ORF belonged.

**ZOT Phylogeny.** The ZOT dataset including outgroups was aligned via MUSCLE (v3.8.31) (Edgar 2004), and the aligned sequences was passed to IQ-TREE (v1.6.1) (Nguyen et al. 2015, Kalyaanamoorthy et al. 2017) with 100 nonparametric bootstraps and the MFP model finder option enabled. Figtree (v1.4.4) was used to visualize the resulting phylogenetic tree (Rambaut 2012).

**T3SS Phylogenies.** For each of the cHGs involved in the inferred transfer one taxa was selected at random and used to query the NCBI non redundant database for high quality matches (cutoff: e-10) via BLASTx (v 2.6.0) (Camacho et al. 2009). These results were classified using the blast option sscinames, and then filtered down to a maximum of 3 members from each genus matched, and no more than 2 taxa of the same species. Furthermore, results were restricted to 20 taxa outside of the *Aeromonas* in order to keep gene trees manageable. These filtered results were then appended to translated cHGs and aligned using MUSCLE (v3.8.31) (Edgar 2004). Aligned sequences were passed to IQ-TREE (v1.6.1) (Nguyen et al. 2015, Kalyaanamoorthy et al. 2017) with 100 nonparametric bootstraps and the MFP model finder option enabled. Figtree (v1.4.4) was used to visualize the resulting phylogenetic tree (Rambaut 2012).

**T3SS Heatmap.** The heatmap for the T3SS was built using R packages ape (v5.3) (Paradis and Schliep 2018), ggtree (v2.0.4) (Yu et al. 2017), phangorn (v2.5.5) (Schliep 2011), phytools (v0.7.20) (Revell 2012), plotly (v4.9.2.1) (Sievert 2020), heatmaply (v1.1.0) (Galili et al. 2017), corrplot (v0.84) (Wei and Simko 2017), corrgram (v1.13) (Wright 2018), dendextend (v1.13.4) (Galili 2015), vegan (v2.5.6) (Oksanen et al. 2019), and Hmisc (v4.4.0) (Jr et al. 2020).

### Supplementary Tables

**Table S1.** Parameter choices for HGT and HMGT inference.

| **HGT inference Parameter** | **Primary setting** | **Other settings used** |
|---|---|---|
| Duplication, transfer, loss costs | $2,4,1$ | $2,3,1$ |
| HGT event support threshold | 100% | – |
| Donor and recipient mapping threshold | $\geq 51\%$ | $\geq 75\%$ |
| **HMGT inference Parameter** | **Primary setting** | **Other settings used** |
| $\langle x,y,z \rangle$ values | $\langle 3,4,1 \rangle$ | $\langle 2,3,1 \rangle$, $\langle 4,5,1 \rangle$, $\langle 5,6,1 \rangle$, $\langle 2,3,2 \rangle$, $\langle 3,4,2 \rangle$, $\langle 4,5,2 \rangle$, $\langle 5,6,2 \rangle$ |
| Rare gene skipping | Skipped | Not skipped |
| Donor or recipient genome ordering | Donor ordering | Recipient ordering |
| cHG ordering or gene ID ordering | cHG ordering | Gene ID ordering for a subset of HGTs |

**Table S2.** Results of HMGT inference analysis on the *Aeromonas* dataset with a stricter mapping threshold of 75%. Results are shown for default settings of all other HGT and HMGT inference parameters (so $\langle x,y,z \rangle = \langle 3,4,1 \rangle$). The table reports (i) the number of donor-recipient (ordered) pairs that had at least one HMGT, (ii) total number of inferred HMGTs, (iii) total number of detected HGTs present within the inferred HMGTs, and (iv) total number of HGTs detected for the reported donor-recipient pairs. These results are shown separately for within-species, across-species, and internal HMGTs.

| | **Pairs** | **HMGTs** | **HMGT-genes** | **HGTs** |
|---|---|---|---|---|
| **Within-species** | 77 | 185 | 617 | 7175 |
| **Across-species** | 66 | 85 | 299 | 1675 |
| **Internal** | 73 | 165 | 565 | 4857 |

**Table S3.** Results of HMGT inference analysis on the *Aeromonas* dataset with a reduced transfer cost of 3. Results are shown for all $\langle x, y, z \rangle$ parameters settings considered, a transfer cost of 3, and default settings for all other parameters (as summarised in Table S1). For each $\langle x, y, z \rangle$ setting, the table reports (i) the number of donor-recipient (ordered) pairs that had at least one HMGT, (ii) total number of inferred HMGTs, (iii) total number of detected HGTs present within the inferred HMGTs (referred to as HMGT-genes), and (iv) total number of HGTs detected for the reported donor-recipient pairs. These results are shown separately for within-species, across-species, and internal HMGTs.

| | Z = 1 | | | | Z = 2 | | | |
|---|---|---|---|---|---|---|---|---|
| | Within-species | | | | Within-species | | | |
| | **Pairs** | **HMGTs** | **HMGT-genes** | **HGTs** | **Pairs** | **HMGTs** | **HMGT-genes** | **HGTs** |
| **X=2, Y=3** | 622 | 3064 | 6942 | 38528 | 622 | 2983 | 7258 | 38528 |
| **X=3, Y=4** | 166 | 493 | 1733 | 16219 | 166 | 473 | 1909 | 16219 |
| **X=4, Y=5** | 59 | 136 | 631 | 8034 | 59 | 134 | 753 | 8034 |
| **X=5, Y=6** | 19 | 34 | 201 | 4140 | 19 | 33 | 220 | 4140 |

| | Across-species | | | | Across-species | | | |
|---|---|---|---|---|---|---|---|---|
| | **Pairs** | **HMGTs** | **HMGT-genes** | **HGTs** | **Pairs** | **HMGTs** | **HMGT-genes** | **HGTs** |
| **X=2, Y=3** | 600 | 1024 | 2360 | 7236 | 600 | 1010 | 2457 | 7236 |
| **X=3, Y=4** | 151 | 200 | 695 | 3453 | 151 | 198 | 735 | 3453 |
| **X=4, Y=5** | 40 | 46 | 219 | 1376 | 40 | 46 | 225 | 1376 |
| **X=5, Y=6** | 13 | 14 | 82 | 519 | 13 | 14 | 83 | 519 |

| | Internal | | | | Internal | | | |
|---|---|---|---|---|---|---|---|---|
| | **Pairs** | **HMGTs** | **HMGT-genes** | **HGTs** | **Pairs** | **HMGTs** | **HMGT-genes** | **HGTs** |
| **X=2, Y=3** | 1002 | 2428 | 5396 | 29824 | 1002 | 2382 | 5599 | 29824 |
| **X=3, Y=4** | 143 | 334 | 1168 | 8676 | 143 | 324 | 1256 | 8676 |
| **X=4, Y=5** | 38 | 84 | 391 | 4179 | 38 | 82 | 430 | 4179 |
| **X=5, Y=6** | 12 | 25 | 141 | 2385 | 12 | 24 | 167 | 2385 |

**Table S4.** Comparison of HMGT inference results using donor species and recipient species orderings. Results are shown for default settings of all other HGT and HMGT inference parameters (so $\langle x,y,z\rangle = \langle 3,4,1\rangle$). The table reports (i) the number of donor-recipient (ordered) pairs that had at least one HMGT, (ii) total number of inferred HMGTs, (iii) total number of detected HGTs present within the inferred HMGTs, and (iv) total number of HGTs detected for the reported donor-recipient pairs. Results are shown separately for within-species, across-species, and internal HMGTs.

**Within-species**

| | **Pairs** | **HMGTs** | **HMGT-genes** | **HGTs** |
|---|---|---|---|---|
| **Donor species orderings** | 144 | 337 | 1135 | 13329 |
| **Recipient species ordering** | 142 | 334 | 1129 | 13016 |

**Across-species**

| | **Pairs** | **HMGTs** | **HMGT-genes** | **HGTs** |
|---|---|---|---|---|
| **Donor species orderings** | 129 | 163 | 561 | 2786 |
| **Recipient species ordering** | 129 | 163 | 559 | 2781 |

**Internal**

| | **Pairs** | **HMGTs** | **HMGT-genes** | **HGTs** |
|---|---|---|---|---|
| **Donor species orderings** | 141 | 345 | 1190 | 9094 |
| **Recipient species ordering** | 144 | 355 | 1241 | 8928 |

**Table S5.** Results of HMGT inference analysis on the *Aeromonas* dataset without skipping over rare genes. Results are shown for default settings of all other HGT and HMGT inference parameters (so $\langle x,y,z\rangle = \langle 3,4,1\rangle$). The table reports (i) the number of donor-recipient (ordered) pairs that had at least one HMGT, (ii) total number of inferred HMGTs, (iii) total number of detected HGTs present within the inferred HMGTs, and (iv) total number of HGTs detected for the reported donor-recipient pairs. Results are shown separately for within-species, across-species, and internal HMGTs.

| | **Pairs** | **HMGTs** | **HMGT-genes** | **HGTs** |
|---|---|---|---|---|
| **Within-species** | 143 | 333 | 1121 | 13269 |
| **Across-species** | 120 | 155 | 522 | 2706 |
| **Internal** | 141 | 337 | 1161 | 9094 |

**Table S6.** Results of HMGT inference analysis on the *Aeromonas* dataset using specific gene IDs instead of cHGs. Results are shown for four different $\langle x,y,z\rangle$ parameter settings and default settings of all other HGT and HMGT inference parameters, and are based on only the subset of HGTs for which gene IDs could be uniquely determined in both the donor and recipient species. For each $\langle x,y,z\rangle$ parameter setting, the table shows the numbers of donor-recipient pairs and HMGTs inferred using cHGs, using gene IDs, and shared in common between them (i.e., their intersections). Results are shown separately for within-species and across-species HMGTs.

**Within-Species**

| | **Pairs** | | | **HMGTs** | | |
|---|---|---|---|---|---|---|
| **parameters** | **cHG** | **Gene ID** | **Common** | **cHG** | **Gene ID** | **Common** |
| **X=2, Y=3, Z=1** | 569 | 568 | 568 | 2478 | 2490 | 2459 |
| **X=3, Y=4, Z=1** | 143 | 146 | 142 | 336 | 344 | 334 |
| **X=4, Y=5, Z=1** | 40 | 42 | 40 | 71 | 72 | 70 |
| **X=5, Y=6, Z=1** | 11 | 11 | 11 | 15 | 16 | 15 |

**Across-Species**

| | **Pairs** | | | **HMGTs** | | |
|---|---|---|---|---|---|---|
| **parameters** | **cHG** | **Gene ID** | **Common** | **cHG** | **Gene ID** | **Common** |
| **X=2, Y=3, Z=1** | 526 | 546 | 525 | 910 | 942 | 905 |
| **X=3, Y=4, Z=1** | 123 | 130 | 121 | 157 | 169 | 155 |
| **X=4, Y=5, Z=1** | 29 | 37 | 29 | 34 | 42 | 34 |
| **X=5, Y=6, Z=1** | 12 | 13 | 12 | 13 | 14 | 13 |

**Table S7.** COG functional categories. The 25 functional categories listed in this table were used in the functional analysis of *Aeromonas* genes; see, e.g., (Tatusov et al. 2000).

INFORMATION STORAGE AND PROCESSING
- [J] Translation, ribosomal structure and biogenesis
- [A] RNA processing and modification
- [K] Transcription
- [L] Replication, recombination and repair
- [B] Chromatin structure and dynamics

CELLULAR PROCESSES AND SIGNALING
- [D] Cell cycle control, cell division, chromosome partitioning
- [Y] Nuclear structure
- [V] Defense mechanisms
- [T] Signal transduction mechanisms
- [M] Cell wall/membrane/envelope biogenesis
- [N] Cell motility
- [Z] Cytoskeleton
- [W] Extracellular structures
- [U] Intracellular trafficking, secretion, and vesicular transport
- [O] Posttranslational modification, protein turnover, chaperones

METABOLISM
- [C] Energy production and conversion
- [G] Carbohydrate transport and metabolism
- [E] Amino acid transport and metabolism
- [F] Nucleotide transport and metabolism
- [H] Coenzyme transport and metabolism
- [I] Lipid transport and metabolism
- [P] Inorganic ion transport and metabolism
- [Q] Secondary metabolites biosynthesis, transport and catabolism

POORLY CHARACTERIZED
- [R] General function prediction only
- [S] Function unknown

**Table S8.** Results of rare-gene HMGT inference analysis for all 103 *Aeromonas* genomes. Results are shown for all $\langle x,y,z \rangle$ parameters settings considered and default settings for all other parameters. For each $\langle x,y,z \rangle$ setting, the table reports (i) the number of genomes that had at least one rare-gene HMGT, (ii) total number of inferred rare-gene HMGTs, (iii) total number of rare genes present within the inferred rare-gene HMGTs, and (iv) total number of rare genes present in the corresponding genomes.

| Parameters | Genomes | Rare-Gene HMGTs | HMGT-genes | Total Rare Genes |
|---|---|---|---|---|
| **x=2, y=3, z=1** | 103 | 2351 | 7044 | 15965 |
| **x=3, y=4, z=1** | 97 | 778 | 3870 | 15769 |
| **x=4, y=5, z=1** | 86 | 382 | 2657 | 15092 |
| **x=5, y=6, z=1** | 73 | 216 | 1973 | 13657 |
| **x=6, y=7, z=1** | 57 | 132 | 1533 | 11359 |

**Table S9.** Estimated false discovery rates for rare-gene HMGTs for all 103 genomes. For each $\langle x,y,z \rangle$ setting, the table reports (i) the average number of genomes (with at least one rare-gene HMGT) and average number of rare-gene HMGTs inferred, (ii) actual numbers of genomes and rare-gene HMGTs, and (iii) the implied false discovery rates for genomes and HMGTs. Results are averaged across 10 randomised runs.

| | **Number of Genomes** | | | **HMGTs** | | |
|---|---|---|---|---|---|---|
| | **Rand. Avg.** | **Actual Total** | **FDR** | **Rand. Avg.** | **Actual Total** | **FDR** |
| **X=2, Y=3, Z=1** | 94.1 | 103 | 91.36% | 1400.8 | 2351 | 59.58% |
| **X=3, Y=4, Z=1** | 42.1 | 97 | 43.40% | 142 | 778 | 18.25% |
| **X=4, Y=5, Z=1** | 8.3 | 86 | 9.65% | 14.1 | 382 | 3.69% |
| **X=5, Y=6, Z=1** | 1.6 | 73 | 2.19% | 1.7 | 216 | 0.79% |

**Table S10.** Estimated false discovery rates for rare-gene HMGTs for the 40 species with at most 100 rare genes. For each $\langle x,y,z \rangle$ setting, the table reports (i) the average number of genomes (with at least one rare-gene HMGT) and average number of rare-gene HMGTs inferred, (ii) actual numbers of genomes and rare-gene HMGTs, and (iii) the implied false discovery rates for genomes and HMGTs. Results are averaged across 10 randomised runs.

| | **Number of Genomes** | | | **HMGTs** | | |
|---|---|---|---|---|---|---|
| | **Rand. Avg.** | **Actual Total** | **FDR** | **Rand. Avg.** | **Actual Total** | **FDR** |
| **X=2, Y=3, Z=1** | 31.2 | 40 | 78.00% | 85.2 | 365 | 23.34% |
| **X=3, Y=4, Z=1** | 1.4 | 34 | 4.12% | 1.6 | 107 | 1.50% |
| **X=4, Y=5, Z=1** | 0 | 24 | 0.00% | 0 | 54 | 0.00% |
| **X=5, Y=6, Z=1** | 0 | 18 | 0.00% | 0 | 31 | 0.00% |

**Table S11.** Functional annotations for cHGs present at the ZOT integration sites shown in Supplementary Figure S11. Note that this table does not include all cHGs present in all ZOT integration sites, only the ones shown in the figure referenced above.

| cHG Number | Annotation |
|---|---|
| 680 | Hypothetical protein |
| 993 | Phage replication initiation protein |
| 1026 | Hypothetical protein |
| 1615 | Phage replication protein |
| 2236 | Transcriptional regulator, HxlR family |
| 2393 | Hypothetical protein |
| 3252 | Phage replication initiation protein |
| 4422 | ZOT |
| 5067 | Hypothetical protein |
| 5381 | Hypothetical protein |
| 5431 | Hypothetical protein |
| 7033 | Flexible pilin precursor |
| 7596 | Hypothetical protein |
| 8593 | Hypothetical protein |
| 8781 | Permease [major facilitator superfamily] |
| 8792 | Hypothetical protein |
| 8855 | Hypothetical protein |
| 9453 | Hypothetical protein |
| 9603 | Minor coat protein |
| 9751 | Hypothetical protein |
| 10085 | Hypothetical protein |
| 10597 | Phage integrase |
| 11010 | ZOT |
| 11176 | Hypothetical protein |
| 11343 | Hypothetical protein |
| 12470 | Hypothetical protein |
| 13061 | Hypothetical protein |
| 13420 | Hypothetical protein |
| 13508 | Aromatic-L-amino-acid decarboxylase |
| 13676 | YebG |
| 14858 | ZOT |
| 15220 | Phage replication protein |
| 16470 | Hypothetical protein |
| 17532 | Hypothetical protein |
| 18844 | 3-hydroxyacyl-dehydratase, FabA form |
| 19491 | Hypothetical protein |
| 19679 | Hypothetical protein |
| 21236 | Hypothetical protein |
| 21420 | Phage protein |
| 22745 | Hypothetical protein |

**Table S12.** Accession numbers for ZOT outgroups.

| Datsaset | Species Name | Protein Accession Number |
|---|---|---|
| ZOT | Pseudomonas stutzeri | WP_015277793 |
| | Campylobacter sp. FOBRC14 | WP_009650613.1 |
| | Campylobacter concisus | WP_103619050.1 |
| | Campylobacter concisus | WP_103624001.1 |
| | Acinetobacter baumannii | WP_032059842.1 |
| | Acinetobacter baumannii | WP_057069658.1 |
| | Vibrio cholerae | WP_071200895.1 |
| | Vibrio cholerae | WP_071919594.1 |
| | Vibrio cholerae | WP_032482437.1 |
| | Pseudomonas stutzeri | WP_015277014.1 |
| | Pseudomonas stutzeri | WP_143510652.1 |
| | Campylobacter sp. FOBRC14 | WP_009650613.1 |
| | Acinetobacter baumannii | WP_032059842.1 |
| | Acinetobacter baumannii | WP_057069658.1 |
| | Campylobacter concisus | WP_103619050.1 |
| | Campylobacter concisus | WP_103624001.1 |
| | Pseudomonas sp. ALS1279 | WP_143510652.1 |
| | Clostridium clostridioforme | SQB14727.1 |
| | Clostridium clostridioforme | SQB14709.1 |
| | Clostridium sp. BL-8 | OOM69526.1 |
| | Clostridium sp. CAG:264 | CCY61697.1 |
| | Clostridium] populeti | SFR97185.1 |
| | Pseudomonas aeruginosa | VFT63508.1 |
| | Burkholderia pseudomallei | VBT23694.1 |
| | Burkholderia pseudomallei | VBI72158.1 |
| | Burkholderia pseudomallei | VCA69511.1 |
| | Pseudomonas aeruginosa | AZP61917.1 |
| | Pseudomonas aeruginosa | AXS73604.1 |
| | Streptococcus pneumoniae | VLS70222.1 |
| | Streptococcus pneumoniae | VRX07622.1 |
| | Streptococcus pneumoniae | VLS32037.1 |
| | Vibrio virus CTXphi | YP_004286239.1 |
| | Vibrio virus CTXphi | AHZ46536.1 |
| | Vibrio virus CTXphi | AHZ46528.1 |
| | Escherichia coli | GCN27534.1 |
| | Escherichia coli | RDS57791.1 |
| | Escherichia coli | RDR14031.1 |

**Table S13.** T3SS HMGT Annotations

| HMGT # | cHG # | Annotation |
|---|---|---|
| 1 | 9080 | type III secretion system inner membrane ring subunit SctD |
| 1 | 11867 | SctC family type III secretion system outer membrane ring subunit AscC |
| 1 | 19570 | Type III secretion system needle filament subunit SctF |
| 2 | 912 | Type III secretion system chaperone AscY |
| 2 | 6421 | LcrR family type III secretion system chaperone |
| 2 | 9090 | LcrG family type III secretion system chaperone AcrG |
| 2 | 9386 | type III secretion system protein AscX |
| 3 | 6421 | LcrR family type III secretion system chaperone |
| 3 | 9090 | LcrG family type III secretion system chaperone AcrG |
| 3 | 21200 | ArcV (Type III secretion cytoplasmic LcrG inhibitor) |
| 3 | 21631 | Type III secretion system translocon subunit AopB |
| 4 | 369 | Putative type III secretion apparatus proteins associated with the locus of enterocyte effacement |
| 4 | 3507 | Type III secretion apparatus protein, HrpE/YscL family |
| 4 | 15341 | Type III secretion inner membrane ring lipoprotein SctJ |
| 5 | 803 | Secretion protein EspA |
| 5 | 19118 | Hypothetical protein |
| 5 | 21179 | Hypothetical protein |
| 6 | 9436 | EscU/YscU/HrcU family type III secretion system export apparatus switch protein |
| 6 | 11240 | Two component system sensor kinase |
| 6 | 11915 | Lytic transglycosylase domain-containing protein |
| 7 | 2121 | Type III secretion system export apparatus subunit SctT |
| 7 | 9436 | EscU/YscU/HrcU family type III secretion system export apparatus switch protein |
| 7 | 17821 | Type III secretion system export apparatus subunit SctS |

Note: Annotations were checked manually against the NCBI non-redundant database. Many are not conclusive annotations.

**Table S14.** Complete listing of *Aeromonas* genomes used, along with statistics on genome completeness, GC content, and genome size.

| Genus | Species | Strain | Completeness % | GC Content | Size (bp) |
|---|---|---|---|---|---|
| *Aeromonas* | *allosaccharophila* | ATCC35942 | 99.73 | 58.09% | 4614823 |
| *Aeromonas* | *allosaccharophila* | BVH88 | 100 | 57.61% | 4775836 |
| *Aeromonas* | *allosaccharophila* | CECT4199T | 100 | 57.43% | 4741632 |
| *Aeromonas* | *austrailiensis* | CECT8023T | 99.73 | 57.17% | 4178588 |
| *Aeromonas* | *bestarium* | CECT4227T | 99.79 | 59.58% | 4769018 |
| *Aeromonas* | *bivalvium* | CECT7113T | 99.63 | 60.39% | 5597214 |
| *Aeromonas* | *caviae* | Ae398 | 99.99 | 60.43% | 4513284 |
| *Aeromonas* | *caviae* | CECT4221 | 99.94 | 60.04% | 4655854 |
| *Aeromonas* | *caviae* | CECT838T | 99.02 | 60.66% | 4545980 |
| *Aeromonas* | *caviae* | TCO22 | 99.46 | 60.18% | 4656131 |
| *Aeromonas* | *dhakensis* | 14 | 99.72 | 60.94% | 4751250 |
| *Aeromonas* | *dhakensis* | 116 | 99.99 | 60.95% | 4755006 |
| *Aeromonas* | *dhakensis* | 145 | 99.91 | 60.44% | 4945219 |
| *Aeromonas* | *dhakensis* | 173 | 99.99 | 60.64% | 4866155 |
| *Aeromonas* | *dhakensis* | 187 | 100 | 60.62% | 4863678 |
| *Aeromonas* | *dhakensis* | 259 | 100 | 60.70% | 4779077 |
| *Aeromonas* | *dhakensis* | 277 | 100 | 60.63% | 4869960 |
| *Aeromonas* | *dhakensis* | AAK1 | 99.9 | 60.74% | 4842937 |
| *Aeromonas* | *dhakensis* | BVH43 | 100 | 60.42% | 5061036 |
| *Aeromonas* | *dhakensis* | BVH65 | 99.71 | 60.72% | 4867583 |
| *Aeromonas* | *dhakensis* | BVH68 | 99.73 | 60.60% | 4941988 |
| *Aeromonas* | *dhakensis* | BVH69 | 100 | 60.65% | 4889942 |
| *Aeromonas* | *dhakensis* | BVH70 | 99.25 | 60.67% | 4799484 |
| *Aeromonas* | *dhakensis* | CECT7289T | 99.6 | 60.75% | 4770582 |
| *Aeromonas* | *dhakensis* | CIP107500 | 100 | 60.76% | 4790446 |
| *Aeromonas* | *dhakensis* | MDS8 | 98.61 | 60.55% | 4922521 |
| *Aeromonas* | *dhakensis* | SSU | 100 | 60.22% | 5023408 |
| *Aeromonas* | *diversa* | CECT4254T | 99.63 | 60.49% | 4130233 |
| *Aeromonas* | *encheleia* | CECT4342T | 99.97 | 60.89% | 4547085 |
| *Aeromonas* | *enteropelogenes* | CECT4255T | 99.59 | 59.02% | 4409548 |
| *Aeromonas* | *enteropelogenes* | CECT4487T | 99.66 | 58.55% | 4549339 |
| *Aeromonas* | *eucrenophila* | CECT4224T | 99.86 | 60.09% | 4615738 |
| *Aeromonas* | *fluvialis* | LMG24681T | 99.18 | 57.30% | 3969336 |
| *Aeromonas* | *hydrophila* | 226 | 99.72 | 59.88% | 5196309 |
| *Aeromonas* | *hydrophila* | ARS13114 | 99.45 | 59.73% | 5027747 |
| *Aeromonas* | *hydrophila* | BAQ071013136 | 99.72 | 59.94% | 5051972 |
| *Aeromonas* | *hydrophila* | CECT839T | 99.72 | 60.54% | 4823523 |
| *Aeromonas* | *hydrophila* | ML09119 | 99.45 | 59.82% | 5108242 |
| *Aeromonas* | *hydrophila* | NF1 | 99.38 | 60.07% | 4886716 |
| *Aeromonas* | *hydrophila* | NF2 | 99.58 | 60.27% | 4866961 |

Continued on next page

Table S14 – continued from previous page

| Genus | Species | Strain | Completeness % | GC Content | Size (bp) |
|---|---|---|---|---|---|
| *Aeromonas* | *hydrophila* | PAQ0910141 | 99.38 | 59.70% | 5036249 |
| *Aeromonas* | *hydrophila* | PAQ09101412 | 99.56 | 60.22% | 5072319 |
| *Aeromonas* | *hydrophila* | PAQ09101421 | 99.72 | 60.06% | 4876716 |
| *Aeromonas* | *hydrophila* | PAQ0910149 | 98.94 | 57.67% | 4679707 |
| *Aeromonas* | *hydrophila* | ranaeCIP107985 | 99.72 | 60.53% | 4760652 |
| *Aeromonas* | *hydrophila* | SNUFPCA8 | 99.45 | 59.83% | 5051930 |
| *Aeromonas* | *jandaei* | CECT4228T | 99.48 | 57.77% | 4575382 |
| *Aeromonas* | *jandaei* | Ho603 | 99.52 | 57.29% | 4730497 |
| *Aeromonas* | *media* | BAQ071013115 | 99.99 | 61.27% | 4639788 |
| *Aeromonas* | *media* | BAQ071013132 | 99.72 | 60.29% | 4771539 |
| *Aeromonas* | *media* | CECT4232T | 99.9 | 59.91% | 4559195 |
| *Aeromonas* | *media* | WS | 99.22 | 60.37% | 4391358 |
| *Aeromonas* | *molluscorum* | CIP108876T | 98.74 | 58.20% | 4306924 |
| *Aeromonas* | *piscicola* | LMG24783T | 99.68 | 57.98% | 5264316 |
| *Aeromonas* | *popoffii* | CIP105493T | 99.6 | 57.44% | 4841897 |
| *Aeromonas* | *rivuli* | DSM22539T | 99.89 | 59.00% | 4609759 |
| *Aeromonas* | *salmonicida* | 01B526 | 99.73 | 57.38% | 5010358 |
| *Aeromonas* | *salmonicida* | 34mel | 100 | 57.52% | 4846648 |
| *Aeromonas* | *salmonicida* | A449 | 99.73 | 57.23% | 5124548 |
| *Aeromonas* | *salmonicida* | achromogenesCIP104001 | 98.02 | 58.05% | 4697918 |
| *Aeromonas* | *salmonicida* | AS03 | 97.92 | 57.72% | 4527153 |
| *Aeromonas* | *salmonicida* | CIP103209T | 99.73 | 57.54% | 4819215 |
| *Aeromonas* | *salmonicida* | masoucidaCIP103210 | 98.85 | 57.76% | 4613513 |
| *Aeromonas* | *salmonicida* | pectinolyticaCIP107036 | 100 | 57.51% | 4883245 |
| *Aeromonas* | *salmonicida* | smithiaCIP104757 | 99.18 | 57.79% | 4601832 |
| *Aeromonas* | *sanarelli* | LMG24682T | 99.71 | 62.08% | 4256736 |
| *Aeromonas* | *schubertii* | CECT4240T | 99.64 | 60.68% | 4195076 |
| *Aeromonas* | *simiae* | CIP107798T | 98.69 | 60.13% | 4054211 |
| *Aeromonas* | *sobria* | 2.01411E+12 | 99.59 | 56.59% | 4729892 |
| *Aeromonas* | *sobria* | ARS14514 | 100 | 56.43% | 4885468 |
| *Aeromonas* | *sobria* | CECT4245T | 100 | 56.68% | 4761755 |
| *Aeromonas* | *sobria* | JG2080 | 100 | 56.54% | 4773357 |
| *Aeromonas* | *sobria* | PAQ0910145 | 100 | 56.59% | 4754673 |
| *Aeromonas* | *sp* | AMC34 | 99.73 | 57.08% | 4655041 |
| *Aeromonas* | *sp nov* | AH4 | 99.91 | 58.63% | 4955315 |
| *Aeromonas* | *taiwanensis* | LMG24683T | 99.29 | 60.68% | 5167084 |
| *Aeromonas* | *tecta* | CECT7082T | 99.63 | 59.04% | 4834645 |
| *Aeromonas* | *veronii* | 159 | 78.71 | 57.80% | 4591719 |
| *Aeromonas* | *veronii* | ADV102 | 99.86 | 57.24% | 3416923 |
| *Aeromonas* | *veronii* | AER39 | 100 | 57.65% | 4601599 |
| *Aeromonas* | *veronii* | AER397 | 100 | 57.47% | 4494268 |
| *Aeromonas* | *veronii* | AK227 | 99.73 | 57.35% | 4571604 |

Continued on next page

**Table S14 – continued from previous page**

| Genus | Species | Strain | Completeness % | GC Content | Size (bp) |
|---|---|---|---|---|---|
| *Aeromonas* | *veronii* | AK236 | 99.32 | 57.78% | 4474914 |
| *Aeromonas* | *veronii* | AK241 | 100 | 57.83% | 4471456 |
| *Aeromonas* | *veronii* | AMC25 | 99.77 | 57.59% | 4676894 |
| *Aeromonas* | *veronii* | AMC35 | 100 | 57.75% | 4677068 |
| *Aeromonas* | *veronii* | B565 | 100 | 57.31% | 4641702 |
| *Aeromonas* | *veronii* | BAQ116 | 100 | 57.76% | 4627647 |
| *Aeromonas* | *veronii* | BAQ135 | 99.73 | 57.68% | 4651682 |
| *Aeromonas* | *veronii* | BVH37 | 100 | 57.89% | 4699570 |
| *Aeromonas* | *veronii* | BVH46 | 100 | 57.85% | 4542408 |
| *Aeromonas* | *veronii* | BVH47 | 99.98 | 57.82% | 4595123 |
| *Aeromonas* | *veronii* | CECT4486 | 99.35 | 57.90% | 4722418 |
| *Aeromonas* | *veronii* | CECT4902 | 100 | 57.84% | 4484342 |
| *Aeromonas* | *veronii* | CECT7059 | 100 | 57.39% | 4719241 |
| *Aeromonas* | *veronii* | CIP107763 | 99.49 | 57.37% | 4888246 |
| *Aeromonas* | *veronii* | F247 | 100 | 57.88% | 4504696 |
| *Aeromonas* | *veronii* | G3C1 | 100 | 57.82% | 4625604 |
| *Aeromonas* | *veronii* | Hm21 | 99.86 | 57.59% | 4843680 |
| *Aeromonas* | *veronii* | Hm22 | 100 | 57.72% | 4882384 |
| *Aeromonas* | *veronii* | LMG13067 | 99.38 | 57.30% | 5010925 |
| *Aeromonas* | *veronii* | TCO21 | 100 | 57.30% | 4814571 |
| *Aeromonas* | *veronii-bv-sobria* | CECT4257T | 99.54 | 57.72% | 4539049 |

**Table S15.** T3SS Gene Tree Accession Numbers.

| Gene Tree | Accession Number | Taxa Name |
|---|---|---|
| 369 | EAA3660113.1 | Salmonella enterica subsp. Enterica |
| 369 | WP_015870332.1 | Edwardsiella ictaluri |
| 369 | ABC60060.1 | Edwardsiella ictaluri 93-146 |
| 369 | WP_081167840.1 | Edwardsiella ictaluri |
| 369 | WP_052654635.1 | Pandoraea oxalativorans |
| 369 | WP_150811427.1 | Pandoraea sputorum |
| 369 | WP_166440858.1 | Chromobacterium vaccinii |
| 369 | WP_070981138.1 | Chromobacterium vaccinii |
| 369 | SUX56120.1 | Chromobacterium violaceum |
| 369 | WP_175007350.1 | Burkholderia lata |
| 369 | WP_150777811.1 | Pandoraea sputorum |
| 369 | WP_099406162.1 | Chitinimonas sp. BJB300 |
| 369 | WP_128899272.1 | Dyella sp. M7H15-1 |
| 369 | WP_145964029.1 | unclassified Chromobacterium |
| 369 | WP_021478898.1 | Pseudogulbenkiania ferrooxidans |

Continued on next page

Table S15 – continued from previous page

| Gene Tree | Accession Number | Taxa Name |
|---|---|---|
| 369 | WP_133681756.1 | Paludibacterium purpuratum |
| 369 | TDR76589.1 | Paludibacterium purpuratum |
| 803 | WP_021475291.1 | Pseudogulbenkiania ferrooxidans |
| 803 | WP_104946547.1 | Chromobacterium vaccinii |
| 803 | WP_046158913.1 | Chromobacterium violaceum |
| 803 | WP_114061647.1 | Chromobacterium sp. IIBBL 112-1 |
| 803 | WP_012847737.1 | Edwardsiella |
| 803 | WP_034171806.1 | Edwardsiella |
| 803 | WP_015870349.1 | Edwardsiella ictaluri |
| 803 | WP_175007382.1 | Burkholderia lata |
| 803 | EAA3660141.1 | Salmonella enterica subsp. Enterica |
| 803 | WP_099406165.1 | Chitinimonas sp. BJB300 |
| 803 | ERE19474.1 | Pseudogulbenkiania ferrooxidans EGD-HP2 |
| 803 | WP_133408707.1 | Parashewanella sp. MEBiC05444 |
| 803 | WP_144046613.1 | Shewanella sp. YLB-06 |
| 803 | WP_133405760.1 | Parashewanella sp. MEBiC05444 |
| 803 | WP_095572000.1 | Vibrio coralliilyticus |
| 803 | WP_006962244.1 | Vibrio |
| 803 | WP_077751519.1 | Shewanella psychrophila |
| 803 | WP_006081435.1 | Shewanella |
| 803 | WP_121839415.1 | Parashewanella curva |
| 803 | WP_038159332.1 | Vibrio |
| 912 | WP_162119973.1 | Photorhabdus bodei |
| 912 | WP_110091827.1 | Photorhabdus luminescens |
| 912 | AAO18049.1 | Photorhabdus luminescens |
| 912 | ALG80913.1 | Yersinia enterocolitica |
| 912 | WP_011117636.1 | Yersinia enterocolitica |
| 912 | WP_002229791.1 | Yersinia |
| 912 | WP_094949072.1 | Pseudomonas sp. IB20 |
| 912 | WP_171958729.1 | Pseudomonas aeruginosa |
| 912 | WP_050704926.1 | Pseudomonas sp. 250J |
| 912 | WP_034249380.1 | Arsenophonus nasoniae |
| 912 | CBA72164.1 | Arsenophonus nasoniae |
| 912 | WP_067360751.1 | Morganella psychrotolerans |
| 912 | WP_046894405.1 | Morganella morganii |
| 912 | WP_146257307.1 | Morganella morganii |
| 912 | WP_009698254.1 | Vibrio |
| 912 | WP_152163565.1 | Vibrio harveyi |
| 912 | WP_144048642.1 | Shewanella sp. YLB-06 |
| 912 | WP_122067846.1 | Vibrio owensii |
| 912 | WP_149027373.1 | Shewanella psychrophila |
| 912 | WP_160348960.1 | Bordetella sp. 15P40C-2 |

Continued on next page

Table S15 – continued from previous page

| Gene Tree | Accession Number | Taxa Name |
|---|---|---|
| 2121 | EAA3660103.1 | Salmonella enterica subsp. Enterica |
| 2121 | WP_106077201.1 | Chromobacterium amazonense |
| 2121 | WP_043627569.1 | Chromobacterium piscinae |
| 2121 | WP_103321780.1 | Chromobacterium sp. MWU13-2610 |
| 2121 | AAX76924.1 | Edwardsiella tarda |
| 2121 | WP_015683273.1 | Edwardsiella |
| 2121 | GAJ67823.1 | Edwardsiella piscicida |
| 2121 | WP_099406149.1 | Chitinimonas sp. BJB300 |
| 2121 | WP_128899283.1 | Dyella sp. M7H15-1 |
| 2121 | WP_150777819.1 | Pandoraea sputorum |
| 2121 | WP_052654530.1 | Pandoraea oxalativorans |
| 2121 | WP_175007373.1 | Burkholderia lata |
| 2121 | WP_150811476.1 | Pandoraea sputorum |
| 2121 | WP_133681765.1 | Paludibacterium purpuratum |
| 2121 | WP_124320326.1 | Pseudomonas chlororaphis |
| 2121 | WP_150052445.1 | Pseudomonas chlororaphis |
| 2121 | WP_087091830.1 | Pseudomonas |
| 2121 | WP_045363061.1 | Mycoavidus cysteinexigens |
| 2121 | WP_026921000.1 | Glomeribacter sp. 1016415 |
| 2121 | WP_114785683.1 | Vibrio sp. A511 |
| 3507 | EAA3660112.1 | Salmonella enterica subsp. Enterica |
| 3507 | WP_106077194.1 | Chromobacterium amazonense |
| 3507 | WP_071110395.1 | Chromobacterium amazonense |
| 3507 | WP_043627543.1 | Chromobacterium piscinae |
| 3507 | WP_034168272.1 | Edwardsiella tarda |
| 3507 | WP_136499181.1 | Edwardsiella tarda |
| 3507 | WP_069578277.1 | Edwardsiella piscicida |
| 3507 | WP_128899271.1 | Dyella sp. M7H15-1 |
| 3507 | WP_152526838.1 | Pseudogulbenkiania ferrooxidans |
| 3507 | ERE16694.1 | Pseudogulbenkiania ferrooxidans EGD-HP2 |
| 3507 | WP_145964030.1 | unclassified Chromobacterium |
| 3507 | WP_052654537.1 | Pandoraea oxalativorans |
| 3507 | WP_150777812.1 | Pandoraea sputorum |
| 3507 | WP_099406163.1 | Chitinimonas sp. BJB300 |
| 3507 | WP_175007351.1 | Burkholderia lata |
| 3507 | WP_150811428.1 | Pandoraea sputorum |
| 3507 | WP_159084720.1 | Gammaproteobacteria bacterium DM2 |
| 3507 | WP_133681758.1 | Paludibacterium purpuratum |
| 3507 | WP_174107949.1 | Cupriavidus gilardii |
| 3507 | WP_150052436.1 | Pseudomonas chlororaphis |
| 6421 | WP_057534778.1 | Yersinia pseudotuberculosis |
| 6421 | WP_050128479.1 | Yersinia pseudotuberculosis |

Continued on next page

**Table S15 – continued from previous page**

| Gene Tree | Accession Number | Taxa Name |
|---|---|---|
| 6421 | WP_002220918.1 | Yersinia intermedia |
| 6421 | AAA98219.1 | Plasmid pCD1 |
| 6421 | WP_036809076.1 | Photorhabdus luminescens |
| 6421 | WP_133815322.1 | Photorhabdus |
| 6421 | WP_110091829.1 | Photorhabdus luminescens |
| 6421 | WP_100846965.1 | Pseudomonas baetica |
| 6421 | WP_070624115.1 | Pseudomonas sp. HMSC075A08 |
| 6421 | WP_033991535.1 | Pseudomonas aeruginosa |
| 6421 | AAT49564.1 | synthetic construct |
| 6421 | WP_026822312.1 | Arsenophonus nasoniae |
| 6421 | WP_171382447.1 | Vibrio europaeus |
| 6421 | WP_069666603.1 | Vibrio europaeus |
| 6421 | WP_005528931.1 | Vibrio campbellii |
| 6421 | RLS12044.1 | Acinetobacter baumannii |
| 6421 | WP_062714130.1 | Grimontia marina |
| 6421 | WP_169915962.1 | Shewanella psychrophila |
| 6421 | WP_050787846.1 | Photobacterium damselae |
| 6421 | WP_077755400.1 | Shewanella psychrophila |
| 9080 | WP_166301237.1 | Photorhabdus cinerea |
| 9080 | WP_112879025.1 | Photorhabdus sp. S8-52 |
| 9080 | WP_166286438.1 | Photorhabdus stackebrandtii |
| 9080 | WP_126559196.1 | Pseudomonas aeruginosa |
| 9080 | WP_070330389.1 | Pseudomonas aeruginosa |
| 9080 | WP_014603715.1 | Pseudomonas aeruginosa |
| 9080 | WP_050139090.1 | Yersinia enterocolitica |
| 9080 | WP_011117640.1 | Yersinia enterocolitica |
| 9080 | WP_011100773.1 | Yersinia enterocolitica |
| 9080 | WP_026822299.1 | Arsenophonus nasoniae |
| 9090 | WP_002212973.1 | Yersinia intermedia |
| 9090 | WP_005176739.1 | Yersinia enterocolitica |
| 9090 | WP_002428714.1 | Yersinia pestis |
| 9090 | WP_094949069.1 | Pseudomonas sp. IB20 |
| 9090 | WP_090291217.1 | Pseudomonas brenneri |
| 9090 | WP_167360831.1 | Pseudomonas brenneri |
| 9090 | WP_077755399.1 | Shewanella psychrophila |
| 9090 | WP_021323770.1 | Photorhabdus temperata |
| 9090 | WP_046973769.1 | Photorhabdus thracensis |
| 9090 | WP_173292598.1 | Shewanella sp. VB17 |
| 9090 | WP_144048639.1 | Shewanella sp. YLB-06 |
| 9386 | WP_112879040.1 | Photorhabdus |
| 9386 | WP_011147904.1 | Photorhabdus laumondii |
| 9386 | WP_054477407.1 | Photorhabdus heterorhabditis |

Continued on next page

**Table S15 – continued from previous page**

| Gene Tree | Accession Number | Taxa Name |
|---|---|---|
| 9386 | WP_010891214.1 | Yersinia enterocolitica |
| 9386 | WP_002212969.1 | Yersinia |
| 9386 | WP_094949073.1 | Pseudomonas sp. IB20 |
| 9386 | WP_025131090.1 | Pseudomonas sp. PH1b |
| 9386 | WP_019250285.1 | Yersinia pestis |
| 9386 | WP_076963253.1 | Pseudomonas |
| 9386 | AAT51637.1 | synthetic construct |
| 9386 | WP_039983248.1 | Vibrio |
| 9386 | WP_054962587.1 | Vibrio bivalvicida |
| 9386 | WP_004745557.1 | Vibrio tubiashii |
| 9386 | NRB24680.1 | Shewanella sp. |
| 9386 | WP_077755403.1 | Shewanella psychrophila |
| 9386 | WP_147507491.1 | Acinetobacter baumannii |
| 9386 | WP_144048643.1 | Shewanella sp. YLB-06 |
| 9386 | WP_086959416.1 | Photobacterium damselae |
| 9386 | WP_062714136.1 | Grimontia marina |
| 9386 | WP_036766111.1 | Photobacterium damselae |
| 9436 | EAA3660102.1 | Salmonella enterica subsp. Enterica |
| 9436 | WP_166440849.1 | Chromobacterium vaccinii |
| 9436 | WP_043627567.1 | Chromobacterium |
| 9436 | WP_046168633.1 | Chromobacterium vaccinii |
| 9436 | WP_136499174.1 | Edwardsiella tarda |
| 9436 | WP_012847748.1 | Edwardsiella |
| 9436 | WP_069578285.1 | Edwardsiella piscicida |
| 9436 | WP_150777818.1 | Pandoraea sputorum |
| 9436 | WP_065225874.1 | Pandoraea oxalativorans |
| 9436 | WP_099406150.1 | Chitinimonas sp. BJB300 |
| 9436 | VVE85793.1 | Pandoraea sputorum |
| 9436 | WP_128899282.1 | Dyella sp. M7H15-1 |
| 9436 | WP_175007374.1 | Burkholderia lata |
| 9436 | WP_133681763.1 | Paludibacterium purpuratum |
| 9436 | WP_097304486.1 | Pseudomonas chlororaphis |
| 9436 | WP_053280205.1 | Pseudomonas chlororaphis |
| 9436 | WP_006081463.1 | Shewanella baltica |
| 9436 | WP_026921001.1 | Glomeribacter sp. 1016415 |
| 9436 | WP_102681240.1 | Pseudomonas |
| 9436 | WP_011846709.1 | Shewanella baltica |
| 11240 | WP_116541284.1 | Edwardsiella piscicida |
| 11240 | ADM40948.1 | Edwardsiella tarda FL6-60 |
| 11240 | WP_012847750.1 | Edwardsiella |
| 11240 | WP_071110387.1 | Chromobacterium amazonense |
| 11240 | WP_043627566.1 | Chromobacterium piscinae |

Continued on next page

**Table S15 – continued from previous page**

| Gene Tree | Accession Number | Taxa Name |
|---|---|---|
| 11240 | WP_158248699.1 | Chromobacterium sp. MWU13-2610 |
| 11240 | WP_175007378.1 | Burkholderia lata |
| 11240 | VVE55963.1 | Pandoraea sputorum |
| 11240 | WP_150777817.1 | Pandoraea sputorum |
| 11240 | WP_099406152.1 | Chitinimonas sp. BJB300 |
| 11240 | WP_128899281.1 | Dyella sp. M7H15-1 |
| 11240 | WP_143855986.1 | Chitinimonas sp. R3-44 |
| 11240 | WP_137937305.1 | Chitinivorax sp. B |
| 11240 | WP_066494370.1 | Burkholderia sp. BDU8 |
| 11240 | WP_059472984.1 | pseudomallei group |
| 11240 | WP_038802126.1 | Burkholderia oklahomensis |
| 11240 | WP_059934381.1 | pseudomallei group |
| 11240 | WP_106698187.1 | Pseudomonas chlororaphis |
| 11240 | WP_097304479.1 | Pseudomonas chlororaphis |
| 11240 | WP_059646874.1 | pseudomallei group |
| 11867 | WP_015060381.1 | Yersinia enterocolitica |
| 11867 | WP_011100762.1 | Yersinia enterocolitica |
| 11867 | WP_050125522.1 | Yersinia enterocolitica |
| 11867 | WP_132352516.1 | Photorhabdus khanii |
| 11867 | WP_165577067.1 | Photorhabdus khanii |
| 11867 | ERT12562.1 | Photorhabdus temperata J3 |
| 11867 | WP_080960377.1 | Pseudomonas |
| 11867 | WP_080960259.1 | Pseudomonas lundensis |
| 11867 | WP_094987257.1 | Pseudomonas lundensis |
| 11867 | WP_162143409.1 | Arsenophonus nasoniae |
| 11867 | WP_034249377.1 | Arsenophonus nasoniae |
| 11867 | PAV73983.1 | Diploscapter pachys |
| 11867 | WP_095759927.1 | Vibrio sp. V1B |
| 11867 | WP_140392093.1 | Vibrio parahaemolyticus |
| 11867 | WP_079750067.1 | Vibrio parahaemolyticus |
| 11915 | EAA3660101.1 | Salmonella enterica subsp. Enterica |
| 11915 | AAV69422.1 | Edwardsiella tarda |
| 11915 | WP_136499173.1 | Edwardsiella tarda |
| 11915 | WP_012847749.1 | Edwardsiella |
| 11915 | WP_150777862.1 | Pandoraea sputorum |
| 11915 | VVE55973.1 | Pandoraea sputorum |
| 11915 | AOZ52984.1 | Chromobacterium vaccinii |
| 11915 | WP_103318001.1 | Chromobacterium sp. MWU13-2610 |
| 11915 | WP_145929354.1 | Chromobacterium vaccinii |
| 11915 | WP_099406151.1 | Chitinimonas sp. BJB300 |
| 11915 | WP_128899950.1 | Dyella sp. M7H15-1 |
| 11915 | WP_175007375.1 | Burkholderia lata |

Continued on next page

**Table S15 – continued from previous page**

| Gene Tree | Accession Number | Taxa Name |
|---|---|---|
| 11915 | OHC62740.1 | Rhodocyclales bacterium GWA2_65_19 |
| 11915 | WP_163135058.1 | Agarivorans sp. Alg241-V36 |
| 11915 | TXT32073.1 | Rhodocyclaceae bacterium |
| 11915 | WP_107152681.1 | Trinickia symbiotica |
| 11915 | WP_169257082.1 | Azoarcus toluvorans |
| 11915 | HCS89263.1 | Chromatiaceae bacterium |
| 11915 | WP_152784980.1 | Agarivorans sp. B2Z047 |
| 11915 | WP_016403241.1 | Agarivorans albus |
| 15341 | EAA3660114.1 | Salmonella enterica subsp. Enterica |
| 15341 | WP_043627545.1 | Chromobacterium |
| 15341 | WP_106077196.1 | Chromobacterium amazonense |
| 15341 | WP_012847722.1 | Edwardsiella |
| 15341 | WP_034171820.1 | Edwardsiella |
| 15341 | WP_104946557.1 | Chromobacterium vaccinii |
| 15341 | WP_081167843.1 | Edwardsiella ictaluri |
| 15341 | WP_021478897.1 | Pseudogulbenkiania ferrooxidans |
| 15341 | WP_150777810.1 | Pandoraea sputorum |
| 15341 | WP_175007349.1 | Burkholderia lata |
| 15341 | WP_052654538.1 | Pandoraea oxalativorans |
| 15341 | WP_150811426.1 | Pandoraea sputorum |
| 15341 | WP_099406161.1 | Chitinimonas sp. BJB300 |
| 15341 | WP_128899273.1 | Dyella sp. M7H15-1 |
| 15341 | WP_133681754.1 | Paludibacterium purpuratum |
| 15341 | WP_071769270.1 | Burkholderia ubonensis |
| 15341 | WP_060236264.1 | Burkholderia ubonensis |
| 15341 | WP_059934372.1 | pseudomallei group |
| 15341 | WP_059582078.1 | pseudomallei group |
| 15341 | QGY32127.1 | Pantoea cypripedii |
| 17821 | EAA3660104.1 | Salmonella enterica subsp. Enterica |
| 17821 | WP_173669975.1 | Edwardsiella ictaluri |
| 17821 | WP_015870358.1 | Edwardsiella ictaluri |
| 17821 | WP_012847746.1 | Edwardsiella |
| 17821 | WP_114061656.1 | Chromobacterium sp. IIBBL 112-1 |
| 17821 | WP_021475303.1 | Chromobacterium violaceum |
| 17821 | WP_103321779.1 | Chromobacterium sp. MWU13-2610 |
| 17821 | WP_150777820.1 | Pandoraea sputorum |
| 17821 | WP_052654527.1 | Pandoraea oxalativorans |
| 17821 | WP_150811477.1 | Pandoraea sputorum |
| 17821 | WP_128899284.1 | Dyella sp. M7H15-1 |
| 17821 | WP_099406148.1 | Chitinimonas sp. BJB300 |
| 17821 | WP_175007372.1 | Burkholderia lata |
| 17821 | WP_028534046.1 | Paludibacterium yongneupense |

Continued on next page

Table S15 – continued from previous page

| Gene Tree | Accession Number | Taxa Name |
|---|---|---|
| 17821 | WP_143855791.1 | Chitinimonas sp. R3-44 |
| 17821 | WP_066494358.1 | pseudomallei group |
| 17821 | WP_133681767.1 | Paludibacterium purpuratum |
| 17821 | WP_059472977.1 | pseudomallei group |
| 17821 | WP_041419869.1 | Shewanella violacea |
| 17821 | WP_144830894.1 | Cupriavidus gilardii |
| 19118 | WP_015870350.1 | Edwardsiella ictaluri |
| 19118 | WP_081167877.1 | Edwardsiella ictaluri |
| 19118 | ABC60077.1 | Edwardsiella ictaluri 93-146 |
| 19118 | WP_166441285.1 | Chromobacterium vaccinii |
| 19118 | WP_021475292.1 | Pseudogulbenkiania ferrooxidans |
| 19118 | WP_046158914.1 | Chromobacterium violaceum |
| 19118 | WP_104946548.1 | Chromobacterium vaccinii |
| 19118 | EAA3660142.1 | Salmonella enterica subsp. Enterica |
| 19118 | VWD65038.1 | Burkholderia lata |
| 19118 | WP_139022873.1 | Chitinimonas sp. BJB300 |
| 19118 | PHV11477.1 | Chitinimonas sp. BJB300 |
| 19118 | WP_175007383.1 | Burkholderia lata |
| 19118 | WP_128898790.1 | Dyella sp. M7H15-1 |
| 19118 | WP_066494390.1 | Burkholderia sp. BDU8 |
| 19118 | WP_059472996.1 | pseudomallei group |
| 19118 | WP_045984622.1 | Vibrio coralliilyticus |
| 19118 | WP_171749521.1 | Vibrio sp. RE88 |
| 19118 | WP_021458126.1 | Vibrio coralliilyticus |
| 19118 | WP_052044788.1 | Arsenophonus endosymbiont of Nilaparvata lugens |
| 19118 | WP_063655022.1 | Candidatus Arsenophonus triatominarum |
| 21179 | EAA3660149.1 | Salmonella enterica subsp. Enterica |
| 21179 | WP_015870351.1 | Edwardsiella ictaluri |
| 21179 | WP_081167880.1 | Edwardsiella ictaluri |
| 21179 | WP_109745744.1 | Edwardsiella piscicida |
| 21179 | WP_043628922.1 | Chromobacterium piscinae |
| 21179 | WP_071110377.1 | Chromobacterium amazonense |
| 21179 | WP_103321770.1 | Chromobacterium sp. MWU13-2610 |
| 21179 | VVE55427.1 | Pandoraea sputorum |
| 21179 | WP_150777781.1 | Pandoraea sputorum |
| 21179 | WP_052654500.1 | Pandoraea oxalativorans |
| 21179 | WP_021475294.1 | Pseudogulbenkiania ferrooxidans |
| 21179 | WP_175007306.1 | Burkholderia lata |
| 21179 | WP_128898791.1 | Dyella sp. M7H15-1 |
| 21179 | WP_133681780.1 | Paludibacterium purpuratum |
| 21179 | WP_099406141.1 | Chitinimonas sp. BJB300 |
| 21179 | WP_028534044.1 | Paludibacterium yongneupense |

Continued on next page

Table S15 – continued from previous page

| Gene Tree | Accession Number | Taxa Name |
|---|---|---|
| 21179 | WP_084637393.1 | Paludibacterium yongneupense |
| 21179 | WP_143855988.1 | Chitinimonas sp. R3-44 |
| 21179 | WP_124656844.1 | Burkholderia sp. Bp9126 |
| 21179 | WP_052902119.1 | Erwinia iniecta |
| 21631 | WP_036775322.1 | Photorhabdus luminescens |
| 21631 | WP_132352486.1 | Photorhabdus khanii |
| 21631 | WP_023045937.1 | Photorhabdus temperata;Photorhabdus temperata J |
| 21631 | WP_094949066.1 | Pseudomonas sp. IB20 |
| 21631 | CBY78186.1 | Yersinia enterocolitica subsp. palearctica Y11 |
| 21631 | WP_011117632.1 | Yersinia enterocolitica |
| 21631 | WP_050336992.1 | Yersinia enterocolitica |
| 21631 | WP_095187046.1 | Pseudomonas sp. Irchel 3E19 |
| 21631 | SDU94459.1 | Pseudomonas brenneri |
| 21631 | WP_002212985.1 | Yersinia pestiss |
| 21631 | SVJ78119.1 | Klebsiella pneumoniae |
| 21631 | WP_069219307.1 | Vibrio harveyi |
| 21631 | WP_038875749.1 | Vibrio jasicida |
| 21631 | WP_104048123.1 | Vibrio jasicida |

### Supplementary Figures

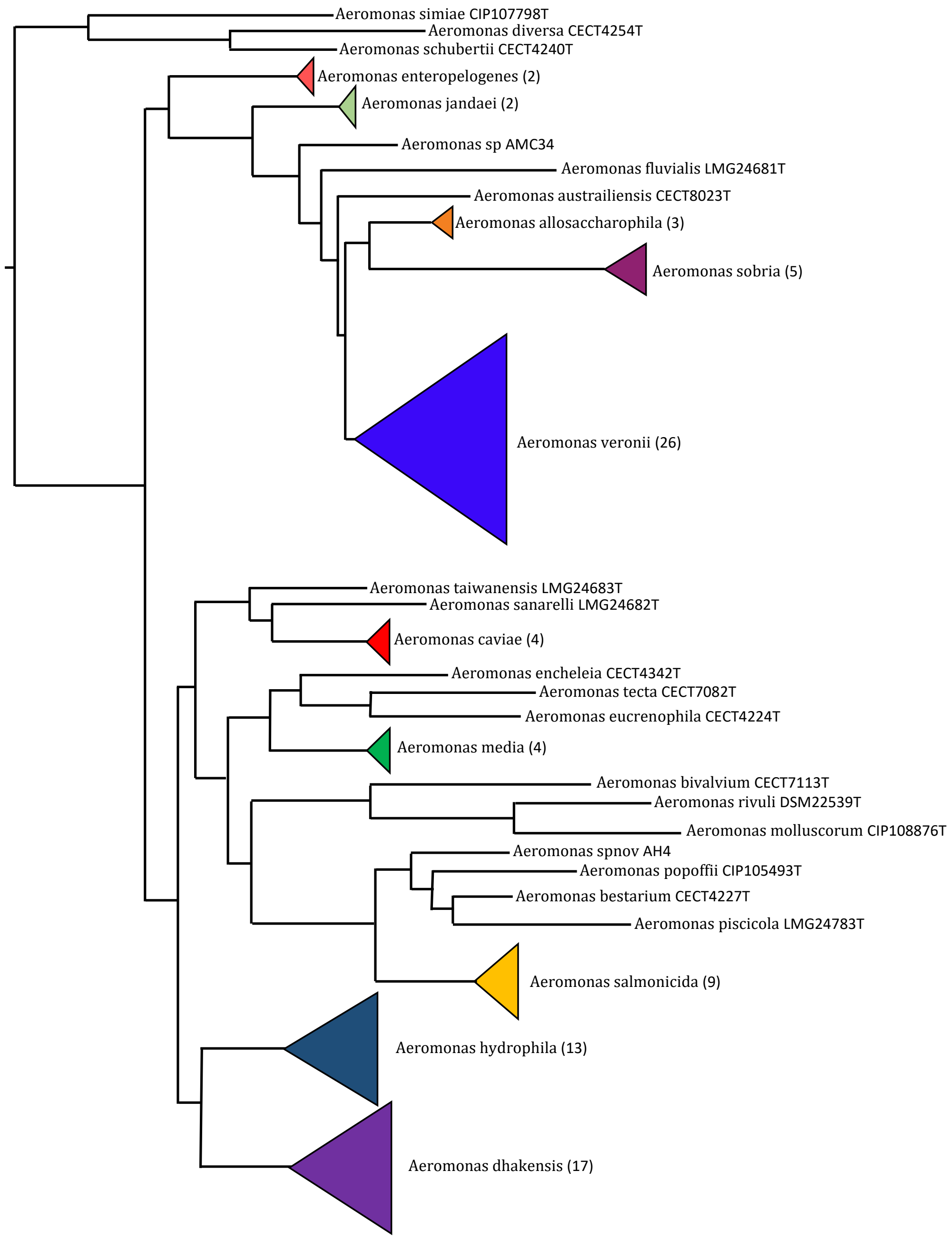


**Figure S1.** Condensed species tree. A condensed view of the species tree used in this study. This rooted tree shows the phylogenetic relationships between the 28 different *Aeromonas* species represented in the analysis. Numbers in parentheses indicate the number of distinct strains (genomes) from that species. The full species tree topology is shown in Figure S2 in the supplement.

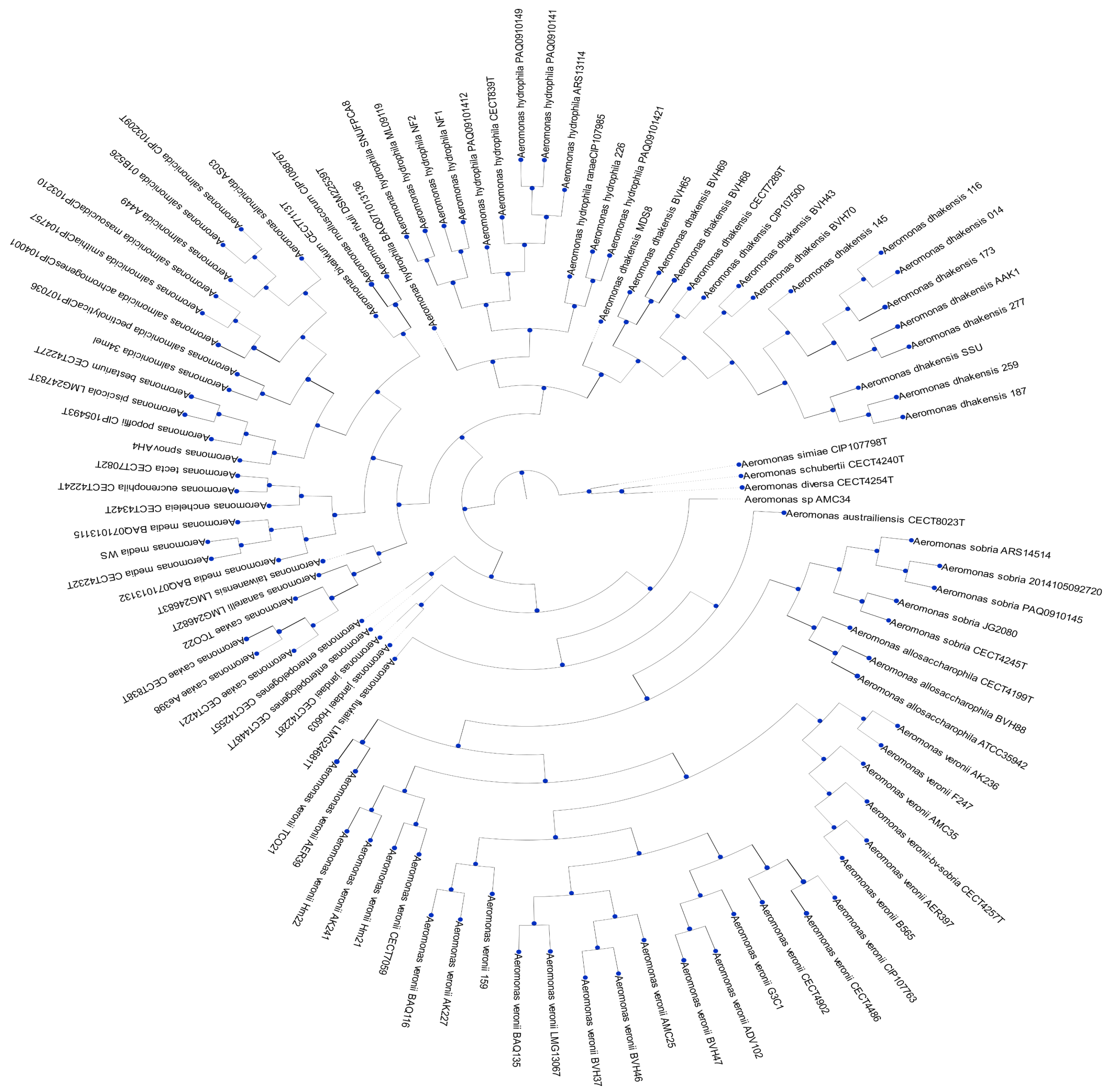


**Figure S2.** Full species tree. A depiction of the phylogenetic relationships (topology only) for the 103 *Aeromonas* genomes used in the analysis. ETE3 (Huerta-Cepas et al. 2016) was used to visualize the phylogeny.

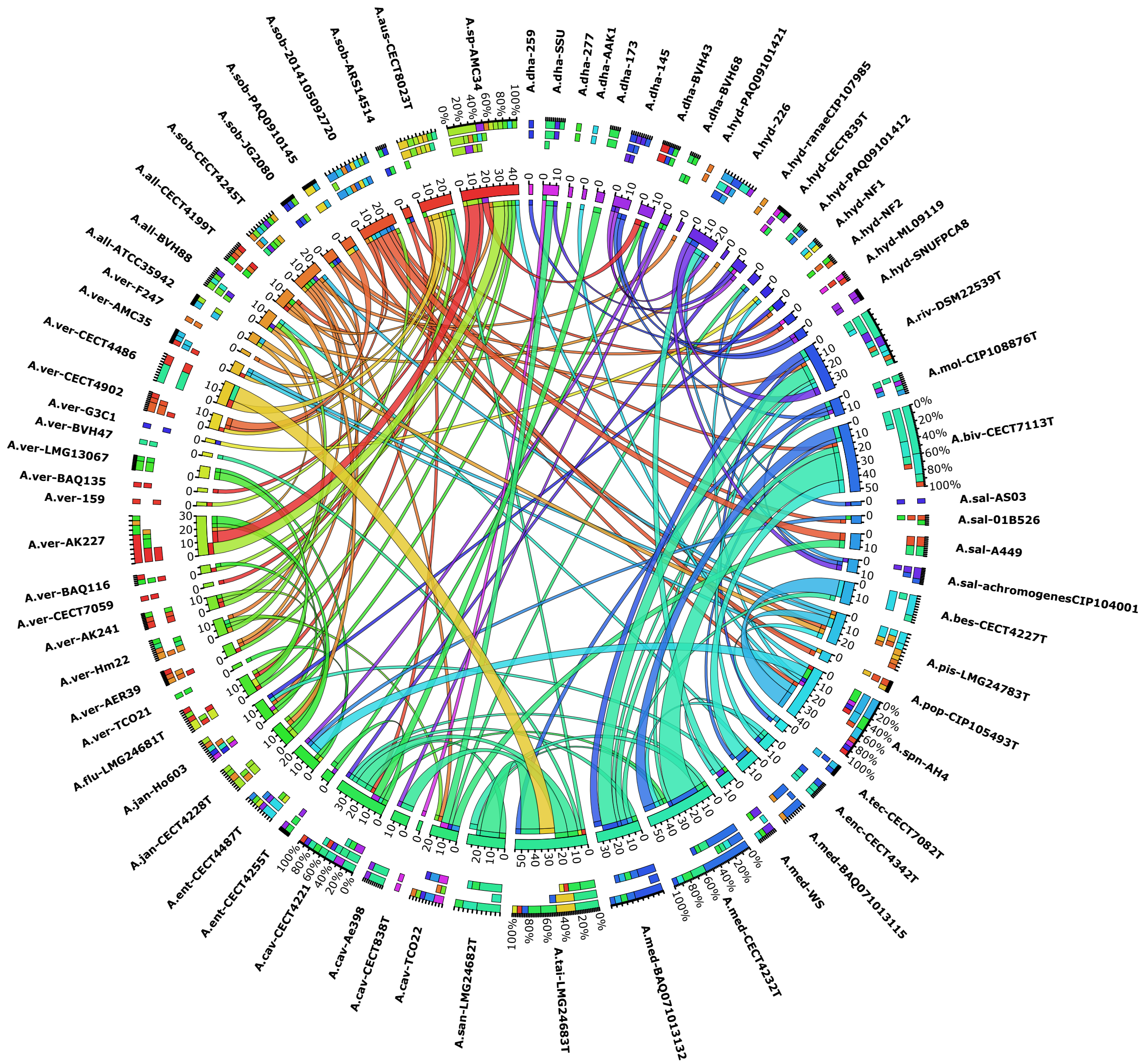


**Figure S3.** Across-species HMGT-genes. Each ribbon connects two *Aeromonas* genomes from different species and corresponds to inferred across-species HMGT-genes between those two genomes. Ribbons are colored according to the color of the donor genome (the color for each genome is shown on the associated segment in the inner ring). The tip of a ribbon at the donor end is colored according to the recipient genome's color. The thickness of a ribbon corresponds to the number of HMGT-genes for that donor-recipient pair, as quantified by the numbers around each segment in the inner ring. For each genome, both incoming (where that genome serves as recipient) and outgoing (where that genome serves as donor) ribbons are shown. The outer ring shows three stacked columns for each genome. Among these three stacked columns, the inner column shows the color distribution of recipients for outgoing ribbons, the middle column shows the color distribution of donors for incoming ribbons, and the outer column shown the combined color distribution for both incoming and outgoing ribbons, for that genome. The figure only includes those *Aeromonas* genomes that served as donor or recipient for at least one across-species HMGT. Only HMGT-genes inferred using default parameters are shown.

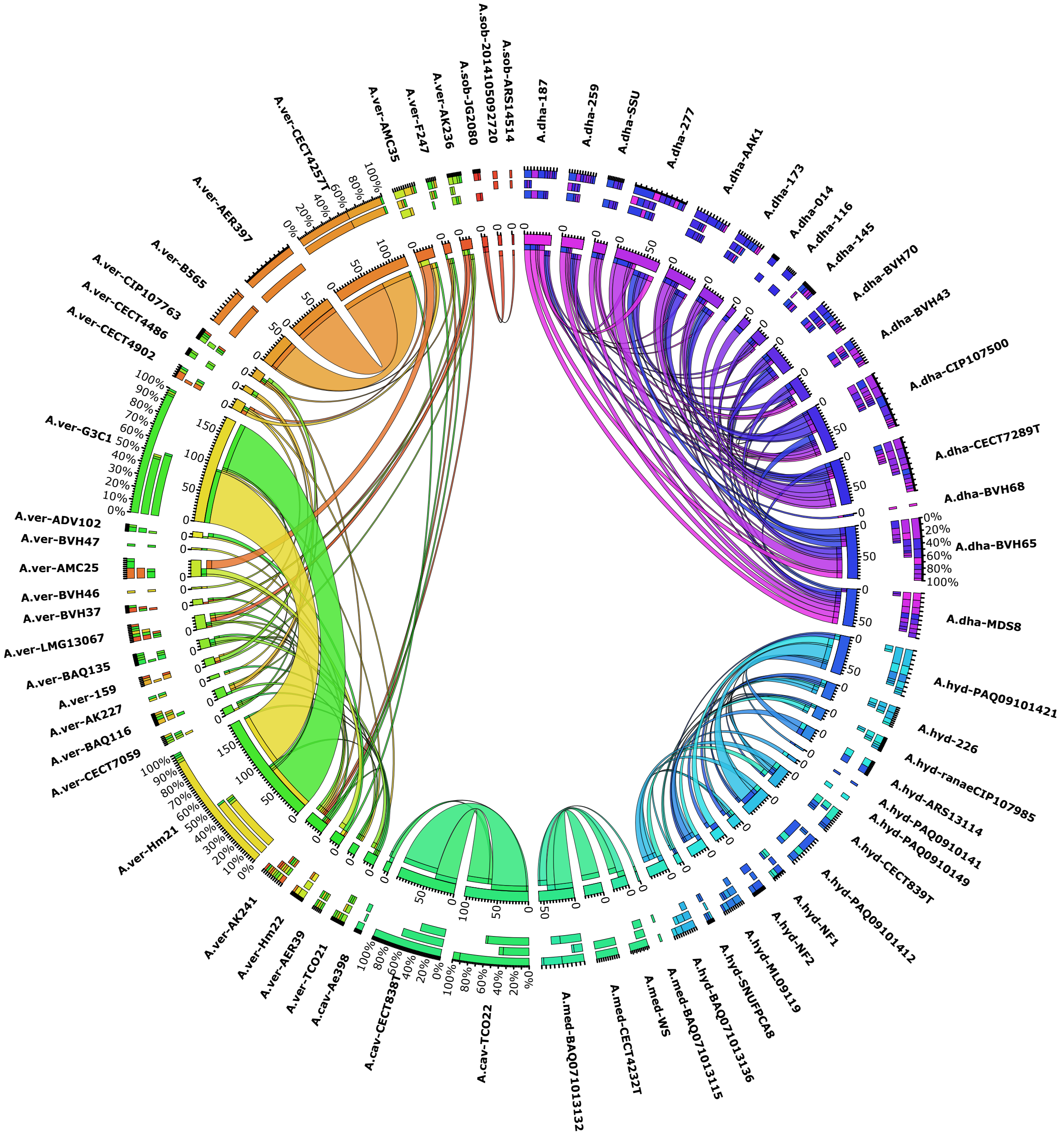


**Figure S4.** Within-species HMGT-genes. Each ribbon connects two *Aeromonas* genomes from the same species and corresponds to inferred within-species HMGT-genes between those two genomes. Ribbons are colored according to the color of the donor genome (the color for each genome is shown on the associated segment in the inner ring). The tip of a ribbon at the donor end is colored according to the recipient genome's color. The thickness of a ribbon corresponds to the number of HMGT-genes for that donor-recipient pair, as quantified by the numbers around each segment in the inner ring. For each genome, both incoming (where that genome serves as recipient) and outgoing (where that genome serves as donor) ribbons are shown. The outer ring shows three stacked columns for each genome. Among these three stacked columns, the inner column shows the color distribution of recipients for outgoing ribbons, the middle column shows the color distribution of donors for incoming ribbons, and the outer column shown the combined color distribution for both incoming and outgoing ribbons, for that genome. The figure only includes those *Aeromonas* genomes that served as donor or recipient for at least one within-species HMGT. Only HMGT-genes inferred using default parameters are shown.

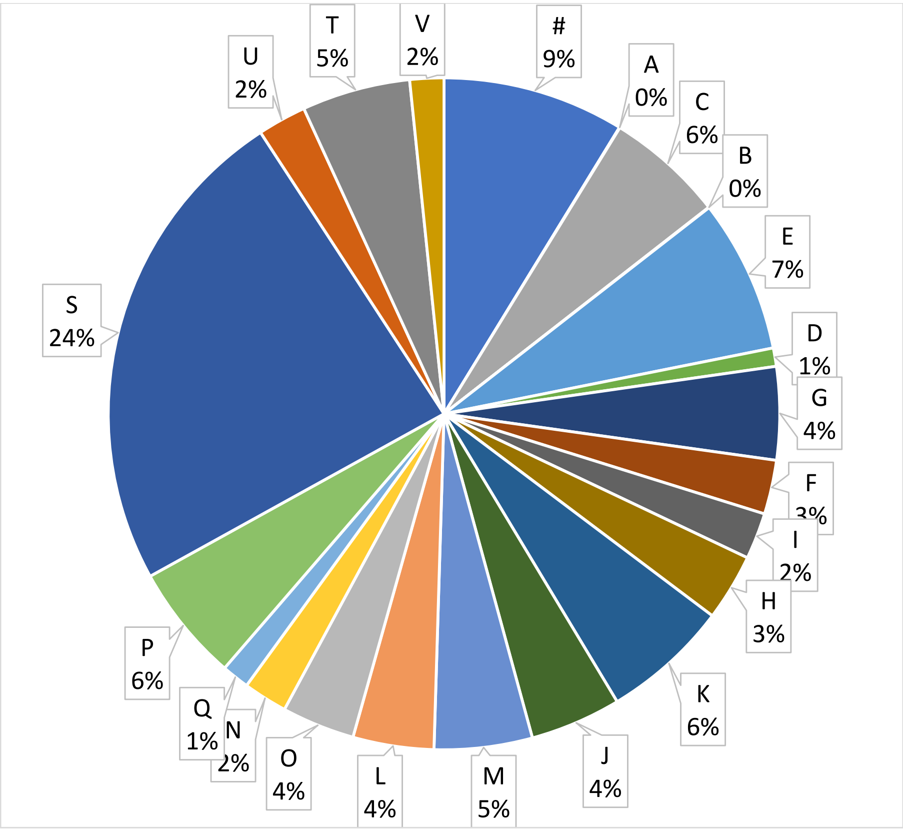


**Figure S5.** Distribution of functional categories for all genes on all genomes. Each letter corresponds to a COG functional category, as detailed in Supplementary Table S7. The "#" character labels those genes for which a COG functional category could not be assigned. COG functional categories "Z", "Y", "W", and "R" are not shown in this pie chart since no gene in any of the *Aeromonas* genomes belonged to those categories.

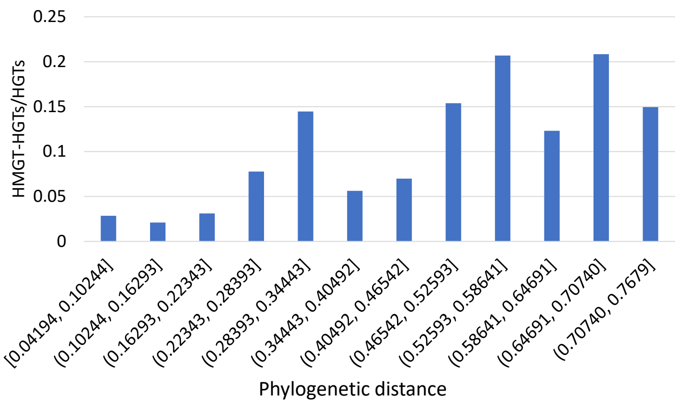


**Figure S6.** Relative HMGT frequencies and phylogenetic distance. This figure shows the ratio of the total number of genes transferred as part of HMGTs (i.e., HMGT-genes) and the total number of detected HGTs for inferred donor-recipient pairs (within- and across-species) corresponding to different phylogenetic distance ranges. Results are shown for donor-recipient pairs, HGTs, and HMGTs inferred using default parameter settings.

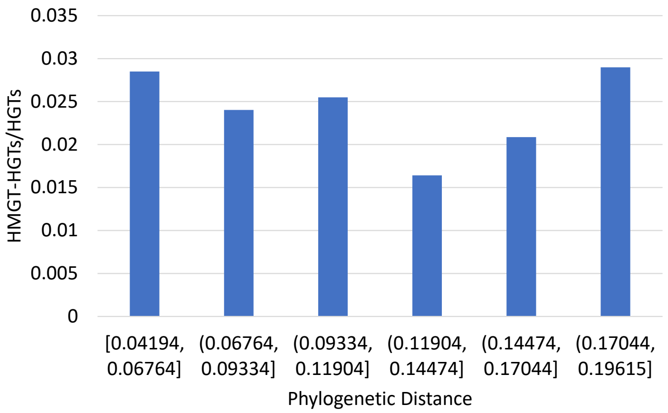


**Figure S7.** Within-species relative HMGT frequencies and phylogenetic distance. This figure shows the ratio of the total number of genes transferred as part of HMGTs (i.e., HMGT-genes) and the total number of detected HGTs for inferred within-species donor-recipient pairs corresponding to different phylogenetic distance ranges. Results are shown for donor-recipient pairs, HGTs, and HMGTs inferred using default parameter settings.

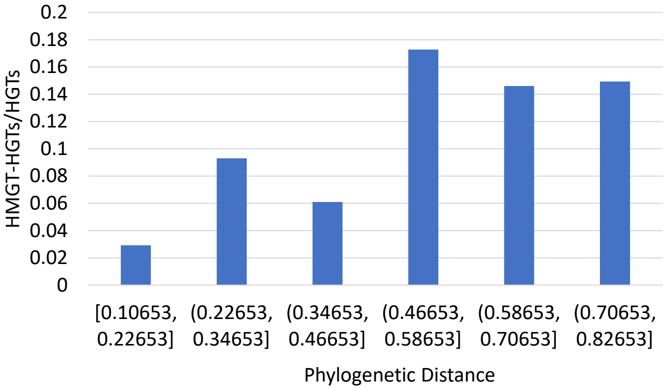


**Figure S8.** Across-species relative HMGT frequencies and phylogenetic distance. This figure shows the ratio of the total number of genes transferred as part of HMGTs (i.e., HMGT-genes) and the total number of detected HGTs for inferred across-species donor-recipient pairs corresponding to different phylogenetic distance ranges. Results are shown for donor-recipient pairs, HGTs, and HMGTs inferred using default parameter settings.

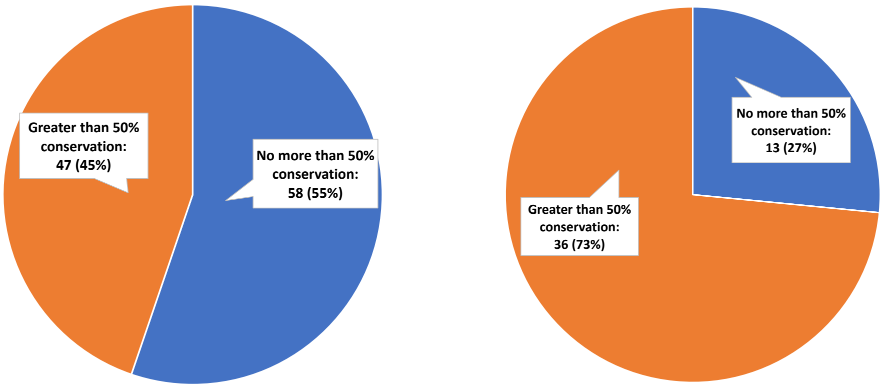


**(a)** Within-species HMGTs **(b)** Across-species HMGTs

**Figure S9.** Functional conservation in HMGTs. The two pie charts show the fraction of HMGTs in which greater than half of their HMGT-genes have conserved COG functions for within-species HMGTs (part (a)) and across-species HMGTs (part (b)). The HMGTs and HMGT-genes used for this analysis were inferred using default parameter settings. Only those HMGTs in which no HMGT-gene was assigned a '#' or [S] functional category were used for this analysis.

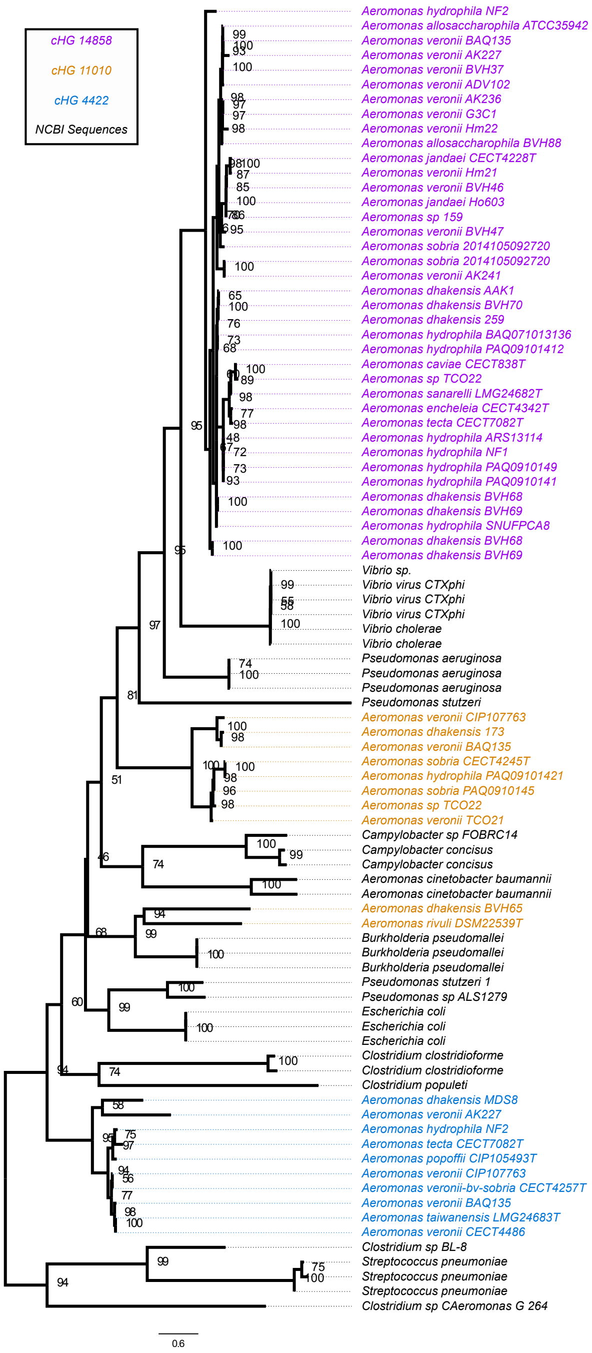


**Figure S10.** ZOT phylogeny with outgroup. IQ-TREE phylogeny of the ZOT toxins from the three cHGs in the Aeromonads and outgroups sampled from NCBI. Taxa are color coded to indicate the group the ZOT sequence is pulled from. Bootstrap values were inferred using IQ-TREE's non-parametric bootstrap option (-b) with 100 replicates.

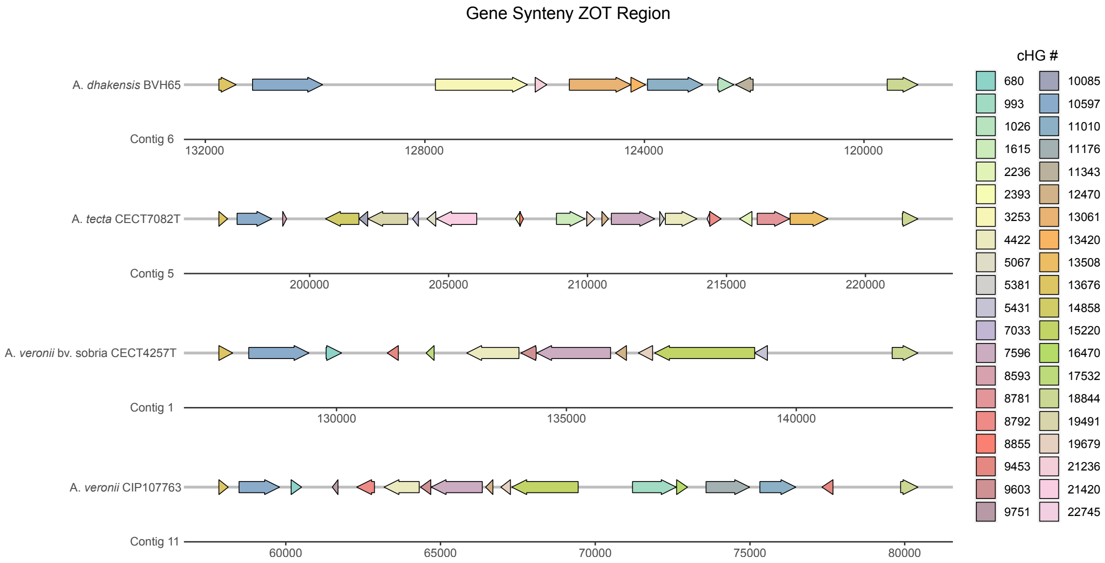


**Figure S11.** Gene synteny plot depicting the diversity of genes and their synteny within the ZOT integration site. Each cHG is colored differently, and a table of cHG annotations is provided in Supplementary Table S11. Arrows depict coding direction and the given numbers correspond to contigs in the draft genomes.

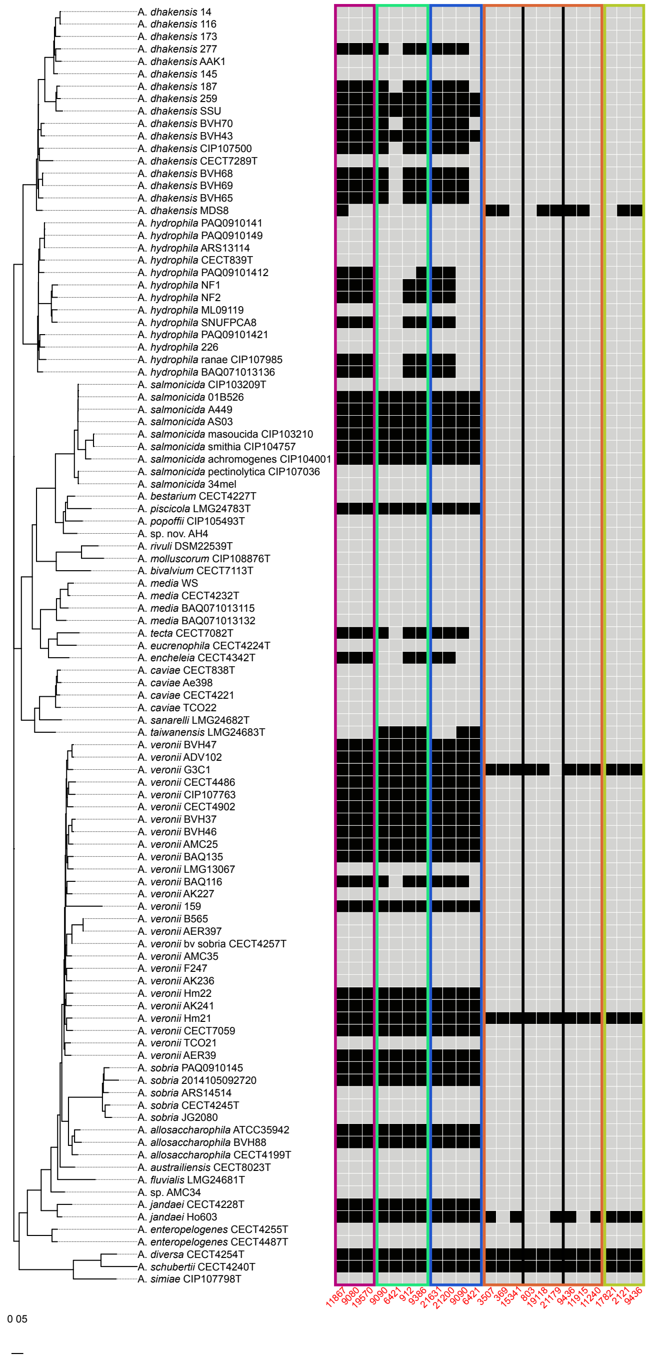


**Figure S12.** Heatmap and unrooted phylogeny showing the presence and absence of genes from inferred T3SS HMGTs. Black boxes indicate presence of the gene, grey indicates absence. Inferred HMGTs are indicated by the colored boxes. The first three boxes (from left to right) are T3SS-1, and the latter 4 are T3SS-2. The orange box separated by black lines are multiple HMGTs that are all transferred between the same two species, whereas all others are from different donors and recipients. Functional annotations for these HMGTs are shown in Supplementary Table S13.

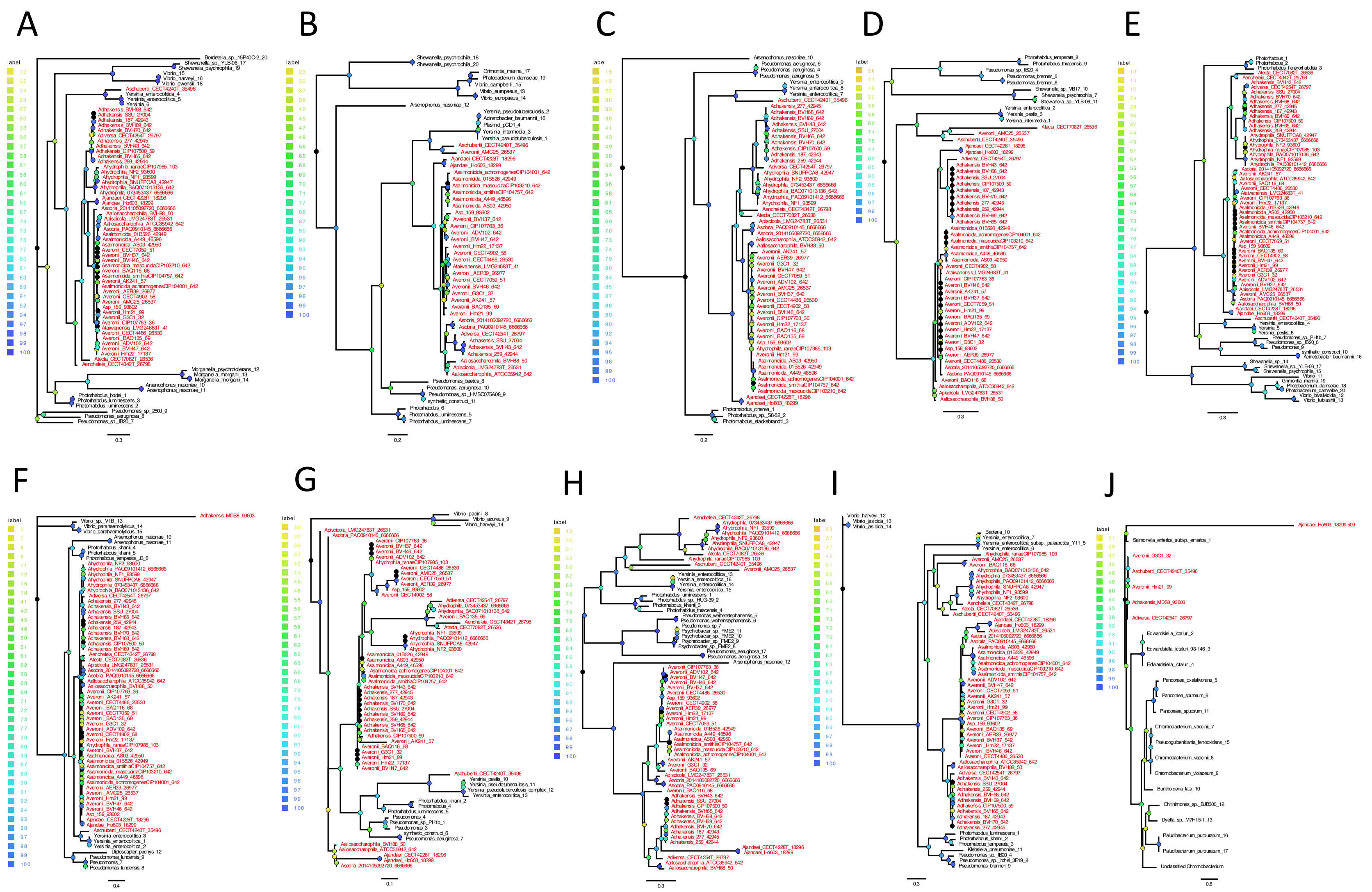


**Figure S13.** See caption on next page. This is page one of a two-page figure.

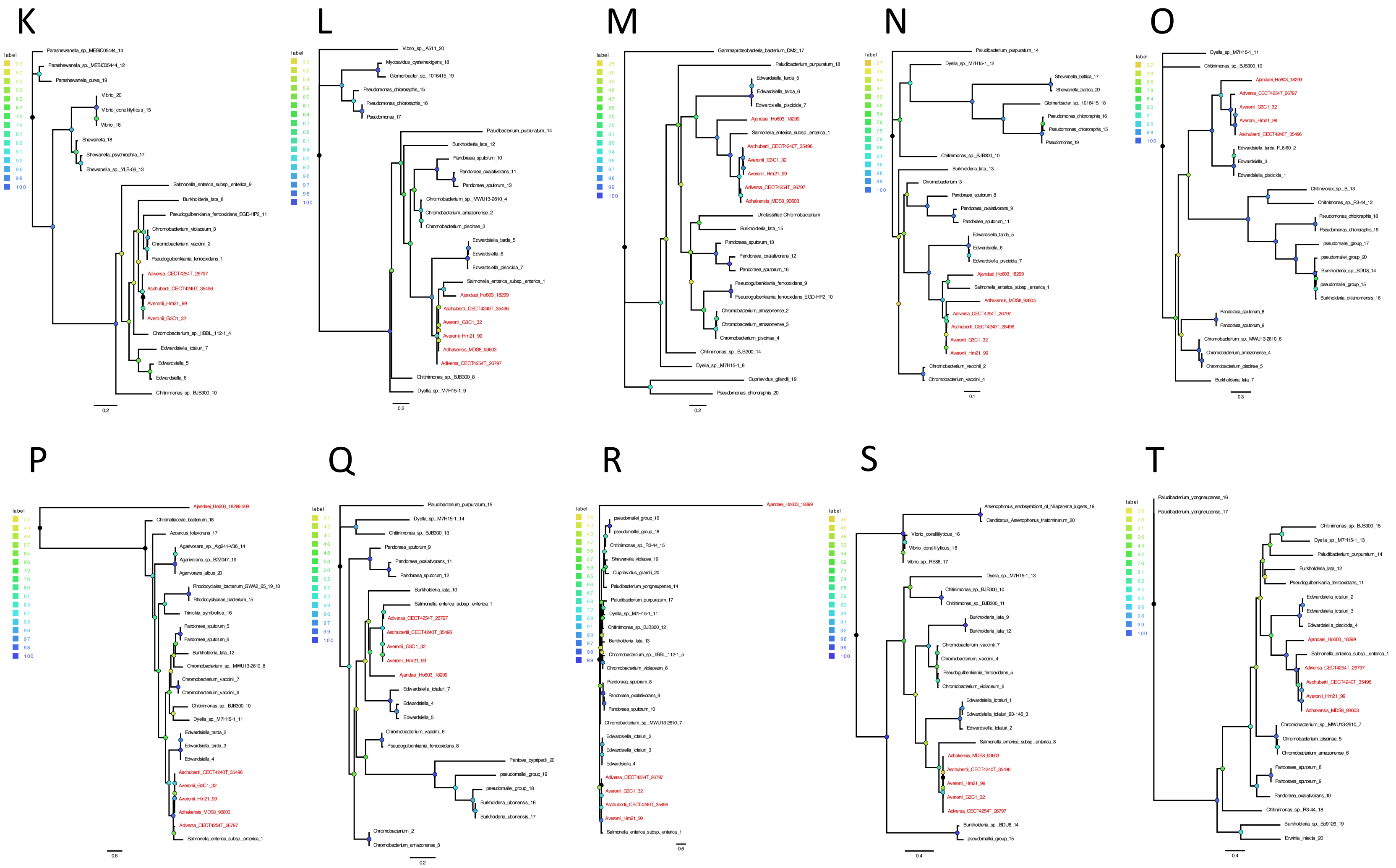

**Figure S13.** Phylogenies of components of Type 3 Secretion Systems. The trees depicted in this two-page figure were calculated from aligned amino acid sequences retrieved from the analyzed *Aeromonas* genomes and from the non-redundant databank; support values were calculated from traditional non parametric bootstrap samples and are color coded according to the insets to the left of each phylogeny. The trees should be considered as unrooted. Panels A-I (shown on the previous page) depict phylogenies of homologs of T3SS-1 components, and Panels J-T depict phylogenies of T3SS-2 components. The correspondence between letter labels and cHGs is as follows. A: 912, B: 6421, C: 9080, D: 9090, E: 9386, F: 11867, G: 19570, H: 21200, I: 21631 (all T3SS-1). J: 369, K: 803, L: 2121, M: 3507, N: 9436, O: 11240, P: 11915, Q: 15341, R: 17821, S: 19118. T: 21179 (all T3SS-2). Accession numbers for all non-*Aeromonas* sequences used in these gene trees (as shown in black) are given in Supplementary Table S15.

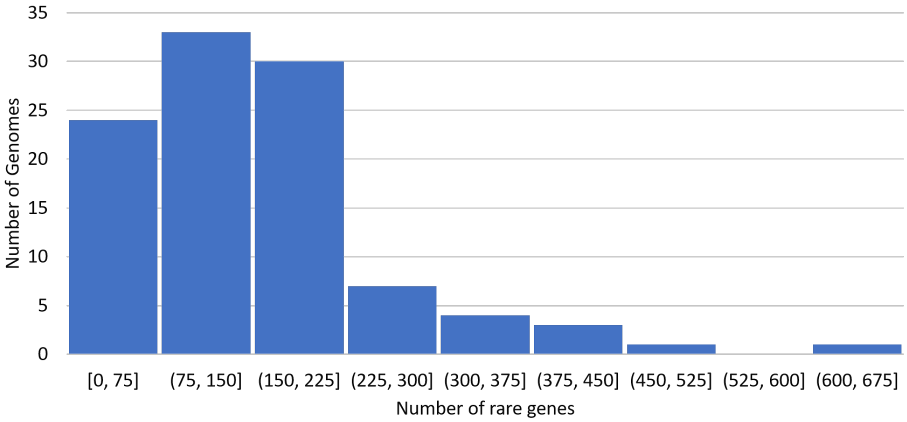


**Figure S14.** Distribution of rare genes across all 103 *Aeromonas* genomes. As the bar chart shows, the vast majority of genomes have no more than 225 rare genes.

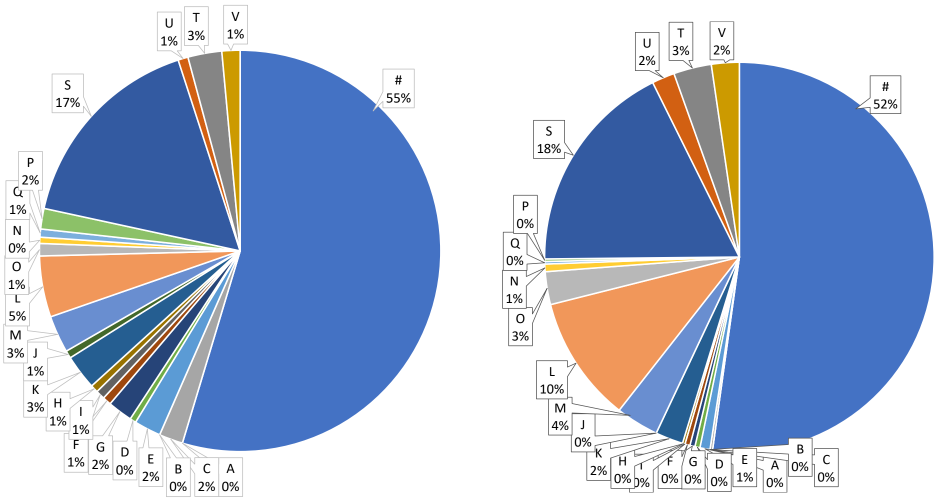


**(a)** Rare genes

**(b)** Rare genes from rare-gene HMGTs

**Figure S15.** The two pie charts show distributions of COG functional categories for all rare genes within the 40 genomes with less than 100 rare genes each (part (a)) and rare genes present in rare-gene HMGTs from those same 40 genomes (part (b)). The rare-gene HMGTs used for this analysis were inferred using the default $\langle x, y, z \rangle$ parameter setting of $\langle 3, 4, 1 \rangle$. Each letter corresponds to a COG functional category, as detailed in Supplementary Table S7. The "#" character labels those genes for which a COG functional category could not be assigned.
